# The Distal-Proximal Relationships Among the Human Moonlighting Proteins: Evolutionary hotspots and Darwinian checkpoints

**DOI:** 10.1101/2023.10.12.562146

**Authors:** Debaleena Nawn, Sk. Sarif Hassan, Moumita Sil, Ankita Ghosh, Arunava Goswami, Pallab Basu, Guy W. Dayhoff, Kenneth Lundstrom, Vladimir N. Uversky

**Affiliations:** Biological Science Division, Indian Statistical Institute, 203 B.T Road, Kolkata, 700108, West Bengal, India; Department of Mathematics, Pingla Thana Mahavidyalaya, Maligram, Paschim Medinipur, West Bengal, India; School of Physics, University of the Witwatersrand, Johannesburg, Braamfontein 2000, South Africa; Woxsen School of Sciences, Woxsen University, Hyderabad - 500 033, Telangana, India; Department of Chemistry, University of South Florida, Tampa, FL 33612, USA; PanTherapeutics, Rte de Lavaux 49, CH1095 Lutry, Switzerland; Department of Molecular Medicine, Morsani College of Medicine, University of South Florida, Tampa, FL 33612, USA

**Keywords:** Moonlighting proteins, Phylogeny, Distal-proximal, Promiscuous proteins, Signature-features

## Abstract

Moonlighting proteins, known for their ability to perform multiple, often unrelated functions within a single polypeptide chain, challenge the traditional “one gene, one protein, one function” paradigm. As organisms evolved, their genomes remained relatively stable in size, but the introduction of post-translational modifications and sub-strategies like protein promiscuity and intrinsic disorder enabled multifunctionality. Enzymes, in particular, exemplify this phenomenon, engaging in unrelated processes alongside their primary catalytic roles. This study employs a systematic, quantitative informatics approach to shed light on human moonlighting protein sequences. Phylogenetic analyses of human moonlighting proteins are presented, elucidating the distal-proximal relationships among these proteins based on sequence-derived quantitative features. The findings unveil the captivating world of human moonlighting proteins, urging further investigations in the emerging field of moonlighting proteomics, with the potential for significant contributions to our understanding of multifunctional proteins and their roles in diverse cellular processes and diseases.

## 1. Introduction

The word moonlighting in English means having an extra job in addition to the canonical task. In the scientific jargon, the meaning for moonlighting proteins is the same for moonlighting proteins whereby the protein along with its mainstream function has an additional function. Because they fulfil numerous independent, frequently unrelated tasks without dividing the functions into discrete protein domains, moonlighting proteins are particularly unique multi-functional proteins [1, 2]. As a result, proteins that emerge from gene fusions and have various functions are disregarded. Proteins that are different splice variants from the same gene are also excluded. Also, both functions should be independent, any changes in one function due to mutation should not affect the other functionality and vice versa. To delineate the phenomenon of moonlighting, Piatigorsky initially coined the term “gene sharing” [3].The term “moonlighting proteins” was first used by Jeffery [4, 5]. The first ever moonlighting function was reported by Piatigorsky and Wistow in 1980s when they reported that certain crystallins, a structural lens protein in the vertebrate eye had other enzymatic functions [6]. In recent years, the conventional paradigm of “one gene encoding one protein with a single function” has yielded to a more nuanced understanding that many proteins exhibit multifunctionality [7]. Amongst these multifunctional proteins, the category of moonlighting proteins stands as a captivating and intricate example. Moonlighting proteins are notable for their capacity to perform two disparate functions within a solitary polypeptide chain. The hypothesis by George Beadle and Edward Tatum stating one gene-one polypeptide-one function cannot be applied in the present scenario [8, 9]. As organisms evolved towards greater complexity, there was not a proportional increase in the size of their genomes. The fundamental genetic code remained unchanged, meaning that the same genes continued to express proteins. However, the emergence of posttranslational modifications (PTMs) in eukaryotes introduced a form of “multi-tasking” into the equation [10]. Two primary strategies for achieving this multi-functionality were protein promiscuity and moonlighting. Moonlighting proteins highlight versatility of evolution and give rise to many interesting evolutionary questions [11, 12, 13, 14]. It is important to note that PTMs themselves played a role in enabling promiscuity and moonlighting, functioning as sub-strategies within these broader concepts [15]. In addition to PTMs, two other notable sub-strategies contributing to this multi-functionality were protein plasticity and intrinsic disorder [15]. Among these sub-strategies, protein promiscuity stood out as a particularly potent mechanism for achieving multi-tasking. This encompassed various forms of promiscuity, such as substrate promiscuity, catalytic promiscuity, and promiscuous protein-protein interactions, all of which played a crucial role in expanding the functional repertoire of proteins [16]. Moonlighting proteins exhibit the remarkable ability to carry out multiple distinct functions autonomously, often without segregating these functions into separate domains within the protein structure [12]. Notably, enzymes serve as striking examples of this phenomenon. In addition to their primary catalytic roles, enzymes are also engaged in entirely unrelated processes, such as autophagy, protein transport, or DNA maintenance [1].

Studies underscore the substantial implications of moonlighting proteins in the context of various diseases and disorders [17, 18, 19, 18]. Notably, while the genomes of many organisms likely harbor a wealth of moonlighting proteins with pivotal roles in pathways and disease etiology, the current roster of confirmed moonlighting proteins remains insufficient to provide a comprehensive vista of the intricate cellular mechanisms that underlie their multifaceted functionality [17].

This quantitative shortfall is primarily attributed to the serendipitous nature by which these proteins’ additional functions are often discovered, emerging incidentally in unrelated experimental contexts [20]. Therefore, the adoption of a systematic quantitative informatics approach holds the potential to deliver significant contributions by identifying hitherto unknown moonlighting proteins and shedding light on the intricate functional attributes inherent to this intriguing class of proteins [21]. This study present a quantitative phylogenetic relationship among various human moonlighting proteins based on quantiative sequence based informatics. This investigation further focused on the complex phenomena between sequence based homology and moonlighting functions of proteins. The research raises thought-provoking questions and proposes hypotheses related to the multifaceted functions of moonlighting proteins. These findings present a promising foundation for forthcoming investigations in the realm of moonlighting proteomics.

## 2. Data acquisition

In the present study, 165 promiscuous proteins/moonlighting proteins of humans were extracted from the database MultitaskProtDBII (Table 1 & Table 12). The list of all these protein sequences including their Uniport ID was supplied as a **Supplementary file-1**.

**Table 1:**
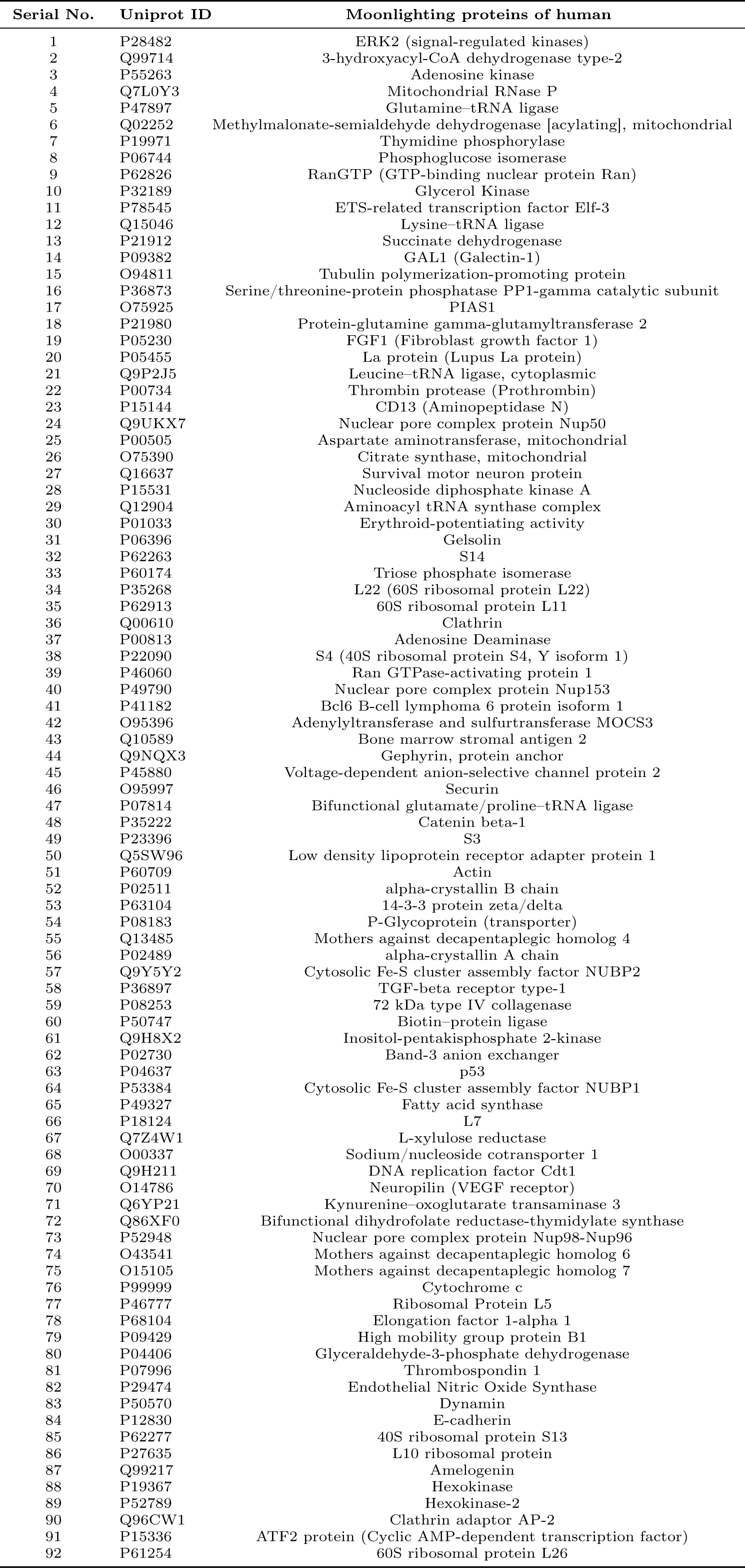
List of Moonlighting proteins in humans with their associated Uniprot ID (Hyperlinked with respective Uniprot webpage)

**Table 2:**
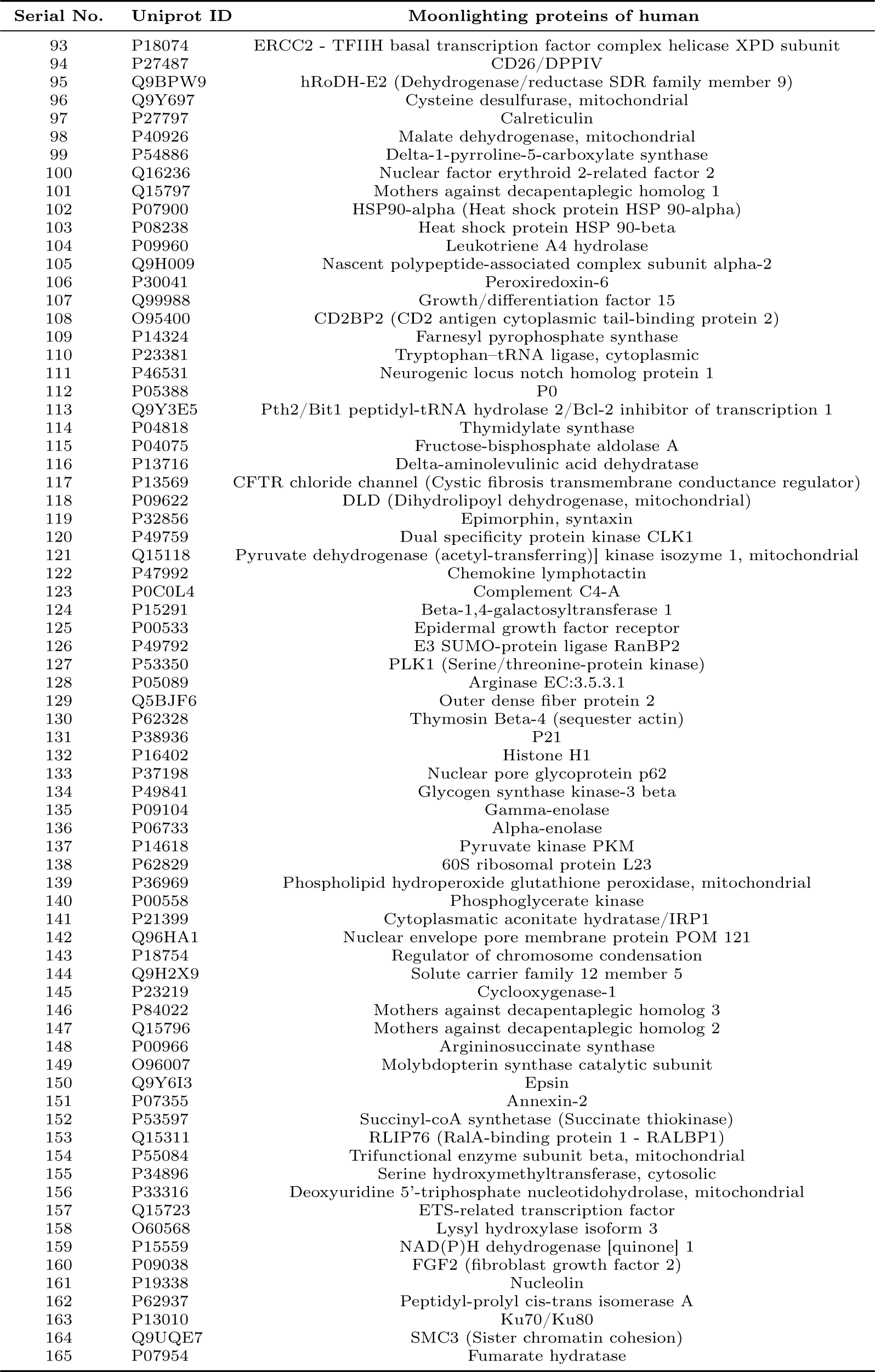
List of Moonlighting proteins in humans with their associated Uniprot ID (Hyperlinked with respective Uniprot webpage)

## 3. Methods

### 3.1. Composition profiler

The Composition Profiler was used to generate an amino acid composition profile of all the human moonlighting proteins analyzed in this study [22]. This set of amino acid sequences was the query set and the ‘Protein Data Bank Select 25’ was the background set. We also generated a composition profile for experimentally validated disordered proteins from the DisProt database. The generated profiles represent plots showing normalized enrichment or depletion of a given residue calculated as 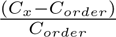, where *C_x_* is the content of a given residue in its query protein, and *C_order_* is the content of the same residue in the PDB Select 25.

### 3.2. Determining amino acid frequency composition

Count of every amino acid in a sequence, termed amino acid frequency, was computed for all 165 moonlighting protein [23, 24, 25]. Due to varied lengths of moonlighting protein sequences, percentage of amino acids in a sequence (obtained from dividing the amino acid frequencies by the length of that sequence and multiplied by 100) is termed the relative frequency of amino acids in that sequence. Relative frequency of 20 amino acids represents a 20 dimensional vector for each protein sequence. Further, 165 moonlighting protein sequences resulted into a 165 dimensional vector for each amino acid. Correlation coefficient matrix (20 × 20) based on all possible pairs of amino acids was computed from the relative frequency vectors of amino acids.

#### 3.2.1. Evaluating Shannon entropy of sequences

Shannon entropy (SE) is a measure of the information content in a system [26]. SE of each moonlighting protein sequence is evaluated by the formula:

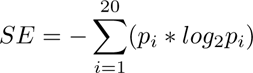

*p_i_* is the count of amino acid *i* in that sequence divided by the length of that sequence [23]. *SE* reflects the degree of randomness in the amino acids count in a given sequence. A higher value of *SE* indicates greater diversity, while a lower value indicates less diversity.

### 3.3. Determining homogeneous poly-string frequency of amino acids

Homogeneous poly-string of length *n* is defined as *n* consecutive occurrence of a particular amino acid [27]. For example, *AAATTAATT* represents one homogeneous poly-string of A with length 3, one homogeneous poly-string of A with length 2, and two homogeneous poly-strings of T with length 2. Note that while counting the number of a homogeneous poly-string of length *n*, only the exclusive/exact occurrence of length *n* would be taken into consideration. The maximum of lengths of homogeneous poly-strings considering all amino acids across all sequences was computed and accordingly, counts of homogeneous poly-strings of all possible lengths (starting from 1 to maximum length) for each amino acid present in a given protein sequence was enumerated.

### 3.4. Evaluating polar, non-polar residue profiles

Every amino acid in a given moonlighting protein sequence was identified as polar(P) or non-polar(Q). Thus, every protein sequence became a binary sequence with two symbols: P and Q. Through this binary P-Q profile, a spatial assembly of polar and non-polar residues over a protein sequence was depicted [28, 29, 30].

#### 3.4.1. Change response sequences based on polar, non-polar residue profiles

There are four possible changes between two consecutive residues of a polar and non-polar profile, namely Polar to Polar (PP), Polar to Non-polar (PN), Non-polar to Non-Polar (NN) and Non-polar to Polar (NP). Such changes were accounted in form of a sequence according to the spatial sequential arrangement of polar, non-polar residues in a given P-Q binary profile. We call this sequence “P-Q Change Response Sequence (*CRS_P Q_*)”. Frequency of each of the four changes was enumerated from binary polar non-polar profile corresponding to each moonlighting protein.

### 3.5. Evaluating acidic, basic, neutral residue profiles

Every amino acid in a given moonlighting protein sequence was identified as acidic (A), basic (B), and neutral (N). Thus, every protein sequence became a ternary valued sequence (A-B-N profiles) with three symbols: A, B, and N.

#### 3.5.1. Change response sequences based on acidic-basic-neutral residue profiles

There are nine possible changes between two consecutive residues of an A-B-N profile, namely Acidic to Acidic (AA), Acidic to Basic (AB), Acidic to Neutral (AN), Basic to Acidic (BA), Basic to Basic (BB), Basic to Neutral (BN), Neutral to Acidic (NA), Neutral to Basic (NB), Neutral to Neutral (NN). Such changes were accounted in form of a sequence according to the spatial sequential arrangement of acidic, basic and neutral residues in a given A-B-N ternary profile. We designate this sequence “A-B-N Change Response Sequence (*CRS_ABN_*)”. Frequency of each of the nine changes was counted for a given ternary A-B-N profile corresponding to each moonlighting protein.

### 3.6. Evaluating intrinsic protein disorder

Predisposition for intrinsic disorder of all the human moonlighting proteins analyzed in this study were determined using a set of commonly used per-residue disorder predictors, such as PONDR® VLS2, PONDR® VL3, PONDR® VLXT, PONDR® FIT, IUPred-Long, IUPred-Short [31, 32, 33, 34, 35, 36]. A web platform called Rapid Intrinsic Disorder Analysis Online (RIDAO) was used to gather results from each predictor in bulk [37]. The percent of predicted intrinsically disorder residues (PPIDR) for each protein was used to classify each protein based on their level of disorder. A residue was considered to be disordered if it had a value of 0.5 or higher. Generally, a PPIDR value of less than 10% is taken to correspond to a highly ordered protein, PPIDR between 10% and 30% is ascribed to moderately disordered protein, and PPIDR greater than 30% corresponds to a highly disordered protein [38, 39]. In addition to PPIDR, mean disorder score (MDS) was calculated for each query protein as a protein length-normalized sum of all the per-residue disorder scores. The per-residue disorder score ranges from 0 to 1, where a score of 0 indicates fully ordered residues and a score of 1 indicates fully disordered residues. Residues with scores above the threshold of 0.5 were considered *disordered residues*. Residues with disorder scores between 0.25 and 0.5 were categorized as *highly flexible*, while those with scores between 0.1 and 0.25 were classified as *moderately flexible* [36]. We also utilized two binary predictors of disorder, the charge-hydropathy (CH) plot and the cumulative distribution function (CDF) (both accessible at Predictor of Natural Disordered Regions), to assess intrinsic disorder at the whole protein level [40, 41]. These binary predictors can be combined to generate CH-CDF plots that distinguish hybrid proteins containing moderate order and disorder from entirely ordered or entirely disordered proteins [42, 43]. Herein, we utilized RIDAO generated CH-CDF plots based on VSL2 CDF scores.

#### 3.6.1. Change response sequences based on intrinsic protein disorder residues

There are sixteen possible changes between two consecutive residues where residues were denoted as disordered (D), highly flexible (HF), moderately flexible (MF) and other (O) in moonlighting protein sequences namely disordered to disordered (D_D), disordered to highly flexible (D_HF), disordered to moderately flexible (D_MF), disordered to other (D_O) and similarly rest twelve (HF_D, HF_HF HF_MF, HF_O, MF_D, MF_HF, MF_MF, MF_O, O_D, O_HF, O_MF, and O_O). Such changes were accounted in the form of a sequence according to the spatial sequential arrangement of D, HF, MF, and O residues for each moonlighting protein sequence. Frequency of each of the sixteen changes was counted from these change response sequences.

### 3.7. Structural and physicochemical features (I-features)

Structural and physicochemical descriptors extracted from sequence data have been widely used to characterize sequences and predict structural, functional, expression and interaction profiles of proteins. iFeature (I-features), a versatile Python-based toolkit was deployed to extract structural and physicochemical properties of the moonlighting proteins [44]. A list of 1182 features which were deployed in the present study was summarized in the following Table 3 [44].

**Table 3:** List of features of dimension 1182 calculated by I-features.

| List of various features (1182 dimensional) calculated by iFeature |  |  |
| --- | --- | --- |
| Descriptor groups | Descriptor | Dimension |
| Amino acid composition | Amino acid composition (AAC) | 20 |
|  | Dipeptide composition (DPC) | 400 |
| Grouped amino acid composition | Grouped Amino Acid Composition (GAAC) | 5 |
|  | Grouped Dipeptide Composition (GDC) | 25 |
| Autocorrelation | Moran Autocorrelation | 240 |
| C/T/D | C/T/D Composition (CTDC) | 39 |
| Conjoint Triad | Conjoint Triad | 343 |
| Quasi-sequence-order | Sequence-Order-Coupling Number (SoC number) | 60 |
| Pseudo-amino acid composition | Pseudo-Amino Acid Composition (PAAC) | 50 |

### 3.8. Formation of distance matrices and dendrograms

Euclidean distance was evaluated between feature vectors of all pairs of moonlighting protein sequences, for each of the following five features: relative frequency of amino acids (dimension 20), relative frequency of changes obtained from polar-nonpolar profiles (dimension 4), relative frequency of changes obtained from acidic-basic-neutral profiles (dimension 9), relative frequency of changes obtained from disordered, highly flexible, moderately flexible, and other residues (dimension 16), structural and physicochemical features (dimension 1182) [45].

Relative frequency was defined as the frequency divided by the length of the sequence and multiplied by 100. Each of the 1182 structural and physicochemical features was normalized in the range of 0 to 100. Each of the five features produces a distance matrix of dimension 165 × 165. For better visualization, distances less than a threshold (empirically chosen separately for each of the distance matrices) were converted to zero. Dendrograms were formed based on Euclidean distances and average linkage. Different color thresholds (empirically chosen) were used in different dendrograms. If the color threshold had the value T, then each group of nodes whose linkage was less than T was assigned a unique color in the dendrogram.

### 3.9. Cumulative clustering of moonlighting proteins

Each of the five distance matrices (as mentioned in subsection 3.7) was normalized by dividing each entry of the matrix by the maximum of that matrix and multiplied by 100. Then all five normalized matrices were added to form a distance matrix for cumulative clustering.

### 3.10. Agglomerated distal-proximal relationships of moonlighting proteins

A moonlighting protein sequence is termed as ‘*outlier*’ with respect to a feature, only if the feature vector of the sequence is distant from most of the feature vectors of the remaining sequences. Set of outlier moonlighting proteins are termed here as *‘distal’* proteins.

A set of moonlighting protein sequences was called as ‘*proximal set*’ with respect to a feature if the sequences belong to the same cluster or immediate adjacent cluster in the dendrogram formed from that feature. Clusters formed from the cumulative clustering were analyzed in detail with respect to each feature to infer how the features individually contributed to the proximal relationship among sequences.

It is worth mentioning that feature extractions and subsequent computations were carried out in *MATLAB* (2023b).

## 4. Results and analyses

### 4.1. Similitude and dissimilitude of moonlighting proteins based on amino acid frequency

#### 4.1.1. Compositional profile of human moonlighting proteins

It is known that the amino acid compositions of ordered and intrinsically disordered proteins are characterized by noticeable difference. In fact, the disordered proteins/regions were shown to be significantly depleted in bulky hydrophobic (I, L, and V) and aromatic amino acid residues (W, Y, F, and H) that are often involved in the formation of the hydrophobic core of a folded globular protein. Disordered proteins/regions also show a low content of C, N, and M residues. These residues, C, W, I, Y, F, L, H, V, N, and M, which are depleted in disordered proteins and regions are defined as order-promoting amino acids. On the other hand, disordered proteins and regions are substantially enriched in disorder-promoting amino acids, such as R, T, D, G, A, K, Q, S, E, and P [46, 47, 35, 48, 22]. These biases in the amino acid composition can be visualized using a web-based tool Composition Profiler for semi-automatic discovery of enrichment or depletion of amino acids in query proteins [22]. Results of the corresponding analysis of 165 human moonlighting proteins are shown in Figure 1. This analysis revealed that out of the ten order-promoting residues six (W, I, Y, H, V, and N) were significantly depleted in the moonlighting proteins, whereas five disorder-promoting residues (R, Q, S, E, and P) were significantly enriched. Curiously, relative to the ordered proteins from the PDB Select 25 set, moonlighting proteins analyzed in this study were significantly enriched in C and L residues while being depleted in D residues.

**Figure 1:**
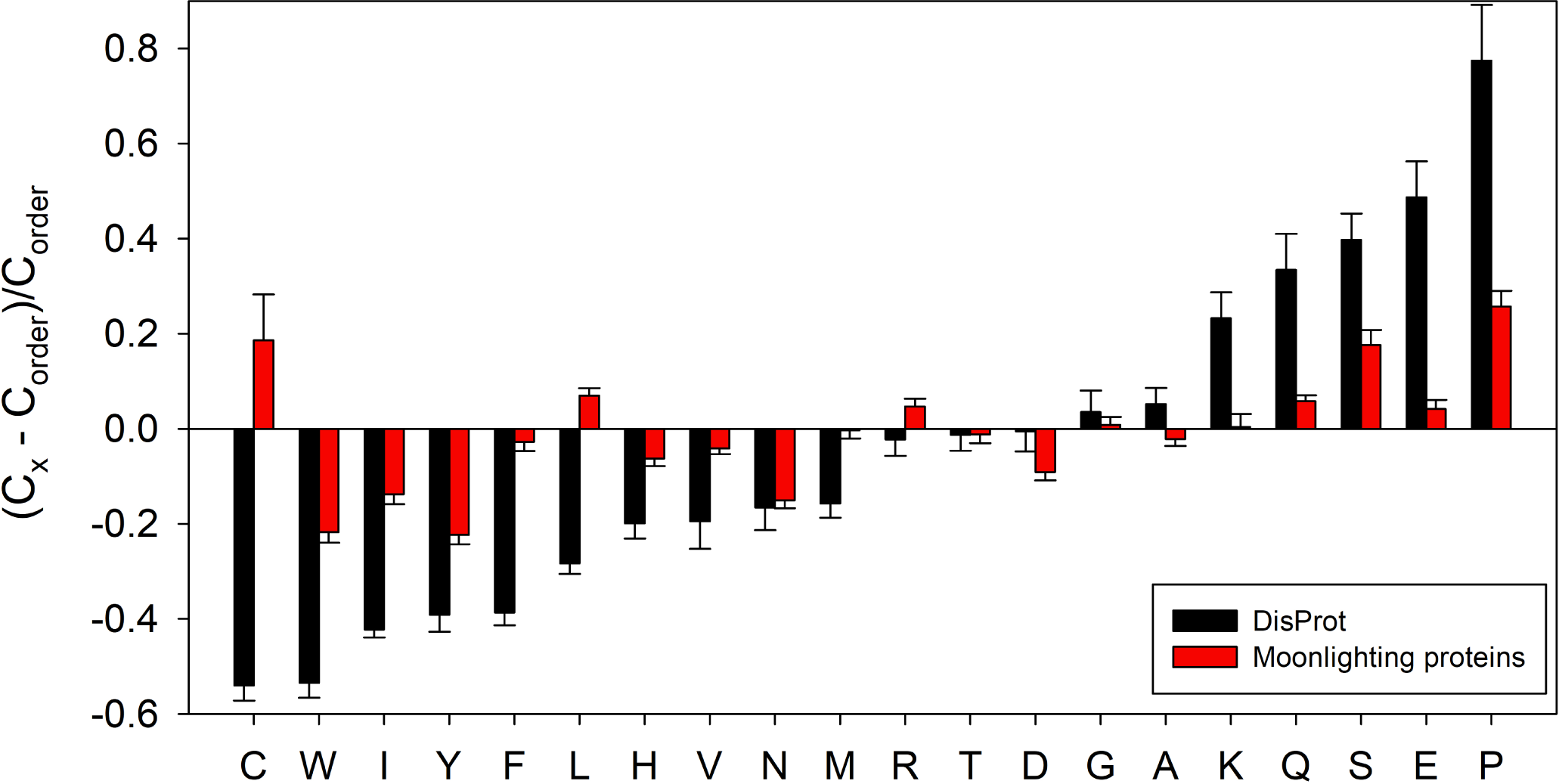
Amino acid composition profile of 165 human moonlighting proteins (red bars). The fractional difference is calculated as 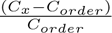, where *C_x_* is the content of a given amino acid in the query set (165 human moonlighting proteins or known intrinsically disordered proteins), and *C_order_* is the content of a given amino acid in the background set (Protein Data Bank Select 25). The amino acid residues are ranked from most order-promoting residue to most disorder-promoting residue. Positive values indicate enrichment, and negative values indicate depletion of a particular amino acid. The composition profile generated for experimentally validated disordered proteins from the DisProt database (black bars) is shown for comparison. In both cases, error bars correspond to standard deviations over 10,000 bootstrap iterations. The composition profile analysis showed that 6 out of 10 order-promoting residues (W, I, Y, H, V, and N) were statistically depleted in the moonlighting proteins (p-value ≤ 0.05), and 5 out of 10 disorder-promoting residue (R, Q, S, E, and P) were significantly enriched in these proteins.

#### 4.1.2. Relative frequency of amino acids and their correlation

It was found that the maximum frequency of Leucine was in 63 sequences followed by alanine (23), glycine (19), lysine (18), serine (13), glutamic acid (12), proline (6), valine (4), aspartic acid (3), threonine (3), and arginine (1) (Numeral inside parentheses denotes the number of moonlighting sequences, where the respective amino acid frequency was found to be maximum). The amino acids that had the minimal frequency along the sequences in the decreasing order were Tryptophan (94) followed by cysteine (42), histidine (9), methionine (9), tyrosine (4), asparagine (2), arginine (1), and phenylalanine (1).

From the relative frequency vector (consisting of 165 elements) of each amino acid, maximum and minimum standard deviation was found for lysine (3.56) and Tryptophan (0.68) respectively (Fig:2). Table 4 shows that only eight amino acids (alanine, glutamic acid, glycine, leucine, lysine, proline, serine, and threonine) were present with more than 15% in twelve sequences.

**Figure 2:**
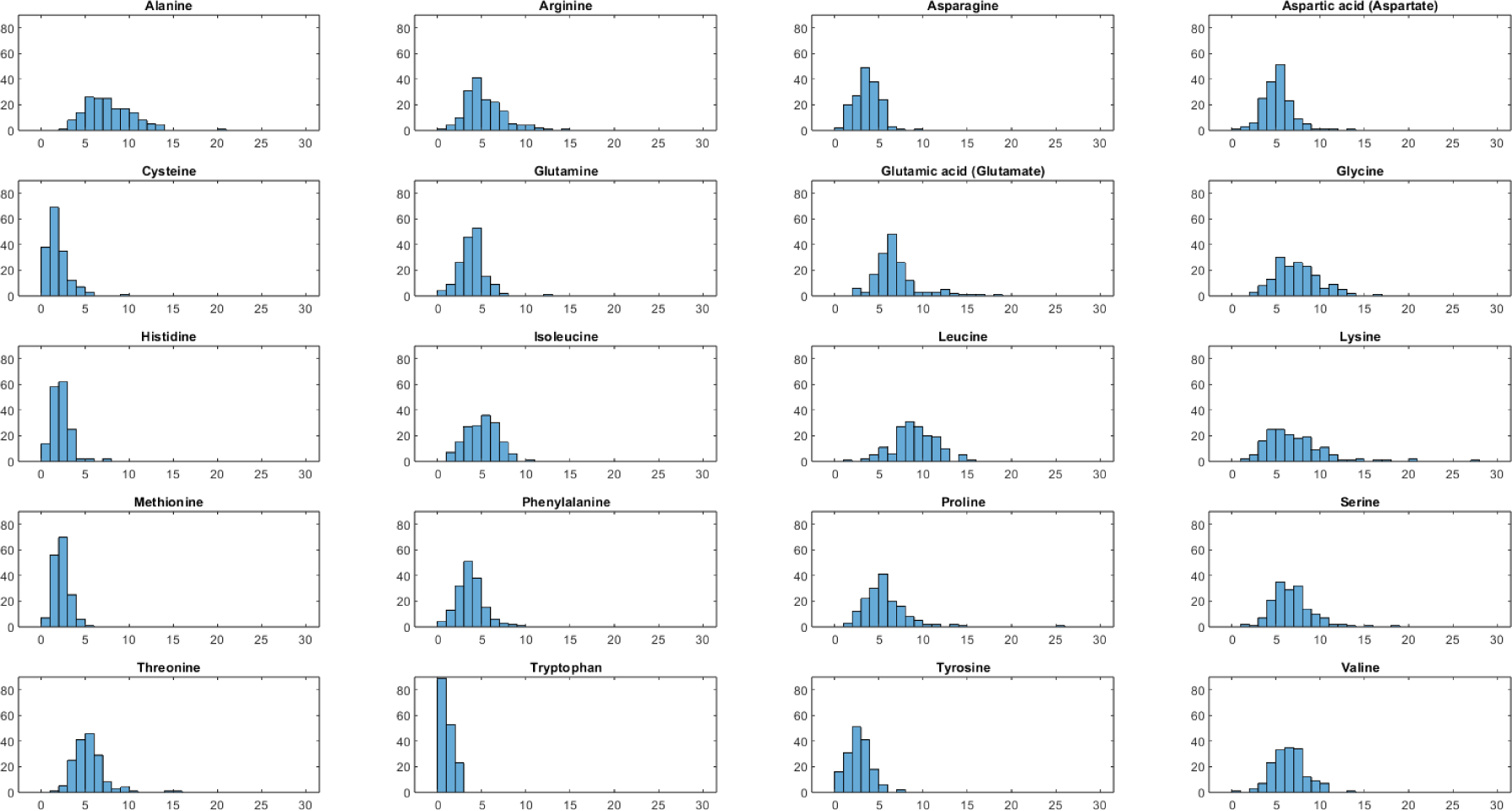
Histogram of relative frequency of each amino acid. X axis denotes percentage and Y axis denotes number of sequences

**Table 4:** Moonlighting sequences with significantly high percentage (≥ 15%) of certain amino acids.

| *Uniprot ID* | alanine | glutamic acid | glycine | leucine | lysine | proline | serine | threonine |
| --- | --- | --- | --- | --- | --- | --- | --- | --- |
| P49790 |  |  |  |  |  |  | 18.1 |  |
| O95396 |  |  |  | 15 |  |  |  |  |
| P99999 |  |  |  |  | 17.14 |  |  |  |
| P09429 |  | 16.74 |  |  | 20 |  |  |  |
| Q99217 |  |  |  |  |  | 25.65 |  |  |
| P61254 |  |  |  |  | 16.55 |  |  |  |
| P62328 |  | 18.18 |  |  | 20.45 |  |  |  |
| P16402 | 20.81 |  |  |  | 27.6 |  |  |  |
| P37198 |  |  |  |  |  |  |  | 15.52 |
| Q96HA1 |  |  |  |  |  |  | 15.21 |  |
| Q15311 |  | 15.57 |  |  |  |  |  |  |
| P09038 |  |  | 16.32 |  |  |  |  |  |

As noticed that the frequency of tryptophan is least in most of the moonlighting protein sequences. One of the reasons for its scarcity being its indole side chain possessing an aromatic bi-nuclear ring structure, unlike the single-ring aromatics found in phenylalanine, tyrosine, and histidine-allowing extensive hydrophobic pockets to be located that facilitate interactions requiring a relatively substantial surface area. Because of these bulky structural characteristics, the biosynthetic pathway of tryptophan stands out as the most energy-demanding among all amino acids, requiring seventy ATP molecules for its production, in contrast to serine, which only requires ten ATP molecules. Tryptophan is coded by only one codon-UGG. Out of sixty-four codons, only one codon is available for its expression. This absence of ambiguity is another reason behind its infrequent appearance [49].

Leucine is a hydrophobic non-polar aliphatic amino acid with a branched side chain. Leucine emerges as a potential amino acid with the greater inclination for *α*-helical structure formation. Among Glu, Ala, Leu, and His, which exhibit the highest occurrences within helical regions, Leu stands out as the predominant residue within the central helical cores of proteins. Therefore a high frequency of Leu suggests that these proteins may have a stable core structure [50]. From the above observations, we can conclude that such a high frequency of amino acids being at the maximal and minimal ends is suggestive of a probable enormous functional diversity. The amino acids present in a higher concentration majorly play a role in key functional regions such as active sites, binding sites, or regions responsible for protein-protein interactions. Also, these amino acids might be preferred due to their ability to form specific secondary structures (e.g., helices or sheets) or specific motifs. Whereas, the amino acids with low frequency conveyed that they are less common and there might be a subset of conserved amino acids that are crucial for the function or structure of the moonlighting protein family.

It was found that absolute values of the Spearman’s rank correlation coefficient for all possible pairs of amino acids were less than 0.5. Hence, no significant positive/negative correlation was obtained in frequency composition of amino acids.

#### 4.1.3. Distance matrix and phylogenetic relationship

Based on a threshold of distance of 8, 126 moonlighting sequences formed 21 clusters as obtained in the dendrogram (Figure 3 (A)). Largest clusters comprised of 35 sequences as marked by brown rectangular box (Figure 3 (A)) (Table 5). Four sequences namely P16402, Q99217, P62328, and P09429 were more than 25 distance away from 153, 136, 43, and 17 sequences, respectively. Among all possible pairwise distances, maximum distance (38.86) was found between P16402 and Q99217. These findings were displayed in form of a distance matrix with a cut off of 25 (Figure 3 (B)).

**Figure 3:**
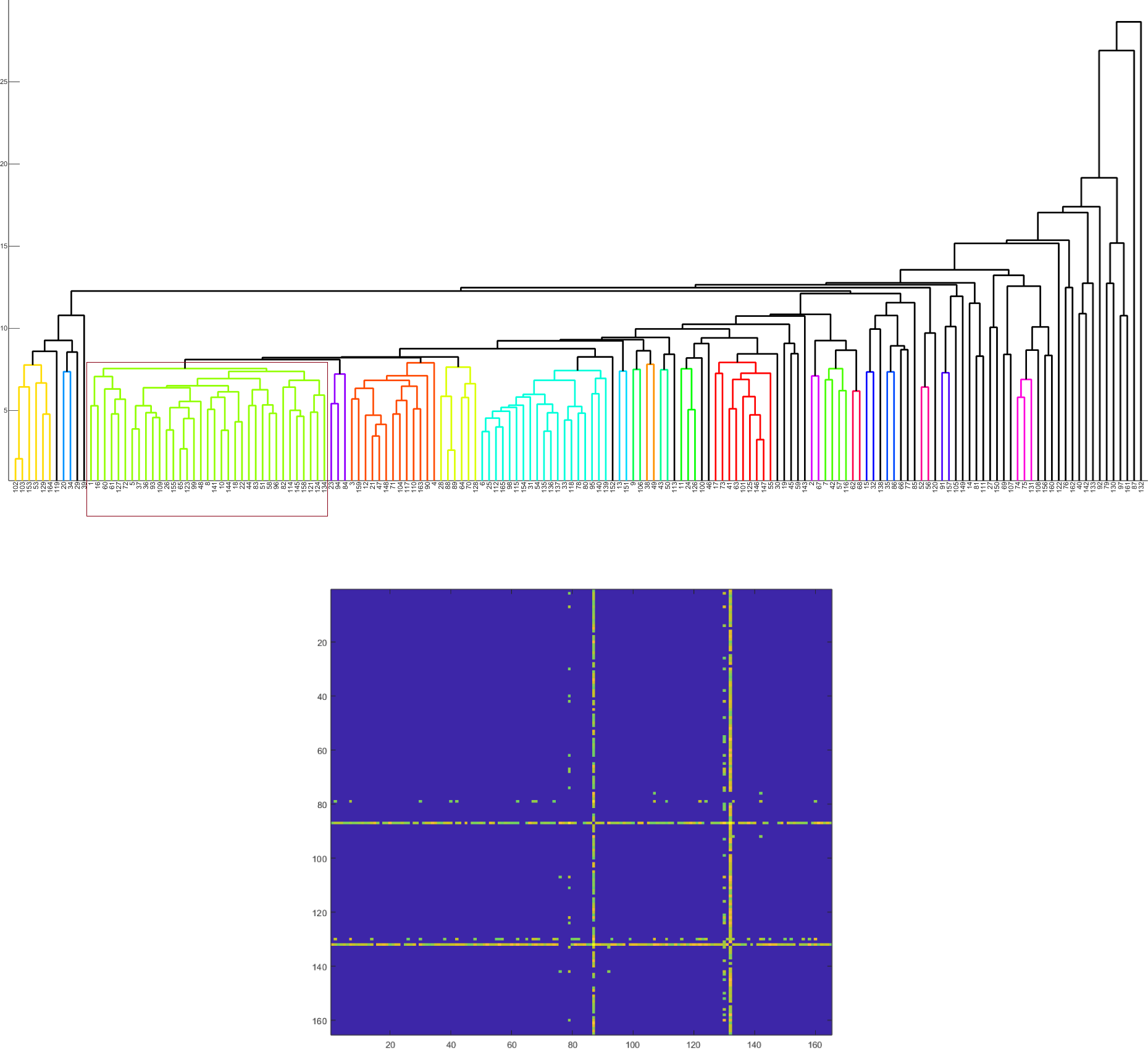
(A): Phylogenetic relationship among the moonlighting proteins based on relative frequency of amino acids. (B): Distance matrix consisting distances between pair of amino acid frequency vectors corresponding to each moonlighting protein (distance less than 25 considered to be zero).

**Table 5:** Clusters derived from relative frequency of amino acids.

| Clusters | Sequences |
| --- | --- |
| Cluster-1 | {102,103,153,53,129,164} |
| Cluster-2 | {20,34} |
| Cluster-3 | {1,16,60,61,127,72,5,37,36,93,109,26,155,65,123,99,48,8,141,10,144,18,22,44,83,51,58,96,82,114,145,158,121,124,134} |
| Cluster-4 | {23,94,84} |
| Cluster-5 | {3,159,12,21,47,148,71,104,117,110,163,90,4} |
| Cluster-6 | {28,88,89,64,70,128} |
| Cluster-7 | {6,25,112,165,98,115,154,31,54,135,136,137,33,118,78,80,95,140,139} |
| Cluster-8 | {13,151} |
| Cluster-9 | {9,106} |
| Cluster-10 | {38,49} |
| Cluster-11 | {43,50} |
| Cluster-12 | {11,24,126} |
| Cluster-13 | {17,73,41,63,101,125,146,147,55} |
| Cluster-14 | {2,67} |
| Cluster-15 | {7,42,57,116} |
| Cluster-16 | {62,68} |
| Cluster-17 | {15,32} |
| Cluster-18 | {35,86} |
| Cluster-19 | {52,56} |
| Cluster-20 | {91,157} |
| Cluster-21 | {74,75,131} |

#### 4.1.4. Shannon entropy of amino acid frequency

Amino acid frequency based Shannon entropy (SE) was calculated for each moonlighting protein sequence. The highest possible SE (4.22) was found for the sequence P84022, whereas the lowest SE (3.19) was obtained for *P* 16402. Sequence P62328 also had a much lower SE (3.47) as obtained in Fig. 4. It was noticed that all four sequences namely P16402, P62328, Q99217, and P09429 which were highlighted in Figure 3 (B) had SEs ranging from 3.19 to 3.75.

**Figure 4:**
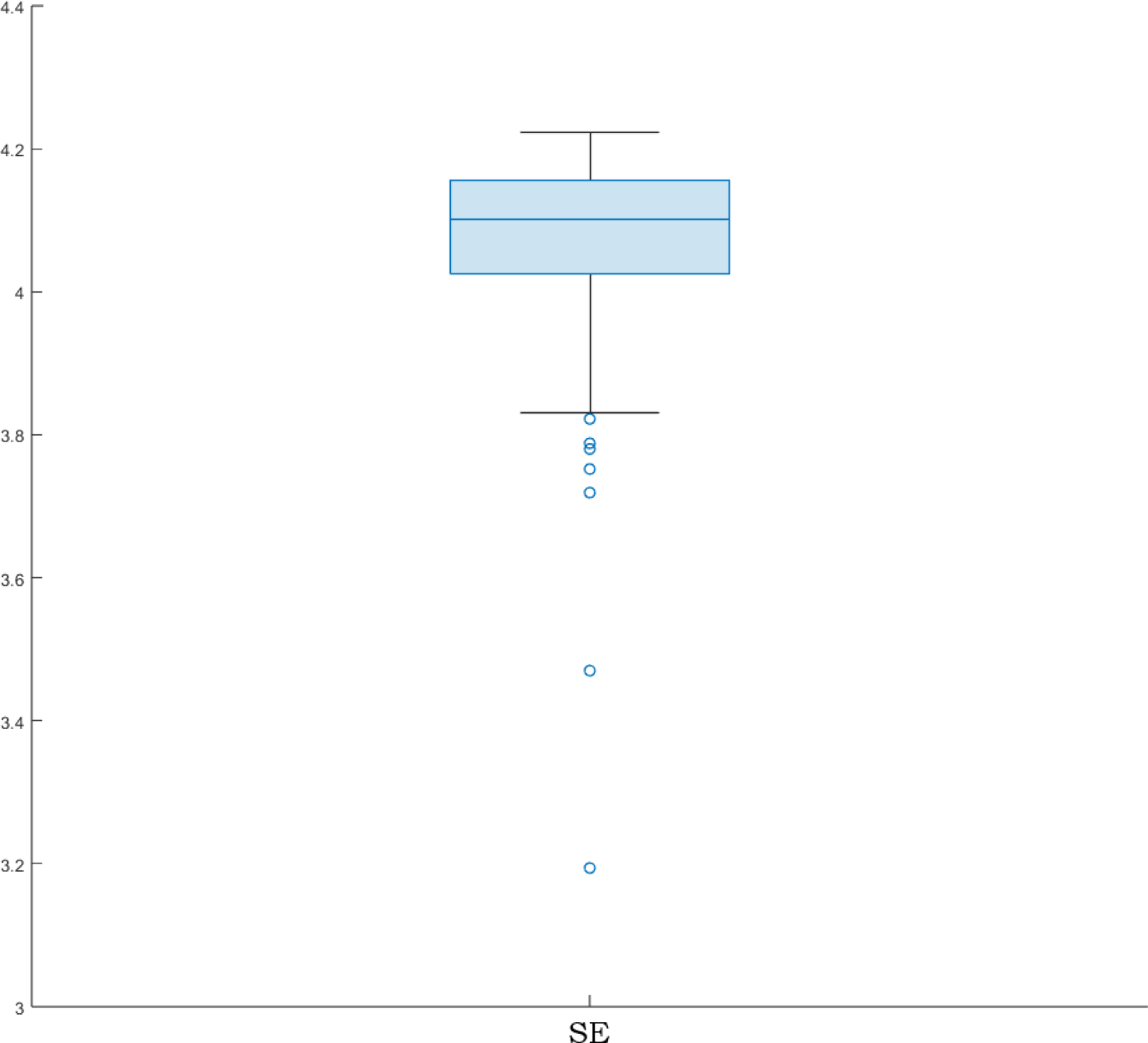
Boxplot of Shannon entropy based on amino acid frequencies derived from respective moonlighting proteins.

The variation in entropy implies the functional versatility and adaptability of moonlighting proteins [26, 51]. Proteins with lower entropy (closer to 3.19) may possess a more specific or constrained set of functions, as their amino acid compositions show less variation [51]. On the other hand, proteins with higher entropy (closer to 4.21) may have more flexible or adaptable functional roles, as their amino acid compositions exhibit greater diversity [52]. Furthermore, it was inferred that the wide range of SEs suggests that moonlighting proteins have undergone evolutionary adaptations to acquire their multiple functions. The different entropy levels likely result from various selective pressures acting on these proteins throughout their evolutionary history [53]. Proteins with higher entropy may have experienced more relaxed selection, allowing for greater divergence in their amino acid compositions and the acquisition of novel functions [54]. Despite the wide range, there may be functional constraints that limit the amino acid composition variability within certain moonlighting proteins [55]. Certain regions or residues may be conserved across the set to maintain essential functional properties required for their moonlighting roles [56]. These conserved features would contribute to the lower end of the entropy range [57].

### 4.2. Homogeneous poly-string frequency of amino acids in moonlighting proteins

The analysis unveiled the maximum length of a homogeneous poly-string to be 14, encompassing all amino acids across the moonlighting protein sequences. This prompted the enumeration of the frequency of homogeneous poly-strings with lengths ranging from 1 to 14 for each of the twenty amino acids, as detailed in **Supplementary file-2**. The cumulative frequency of all homogeneous poly-strings was then calculated for each moonlighting protein and summarized in Table 6. Interestingly, no poly-strings of lengths 8, 11, 12, and 13 were present in any of the 165 moonlighting protein sequences. A single instance of a poly-string length of 14 was observed in the moonlighting sequence P46060, composed of glutamic acid and occurring with a frequency of 1.

**Table 6:**
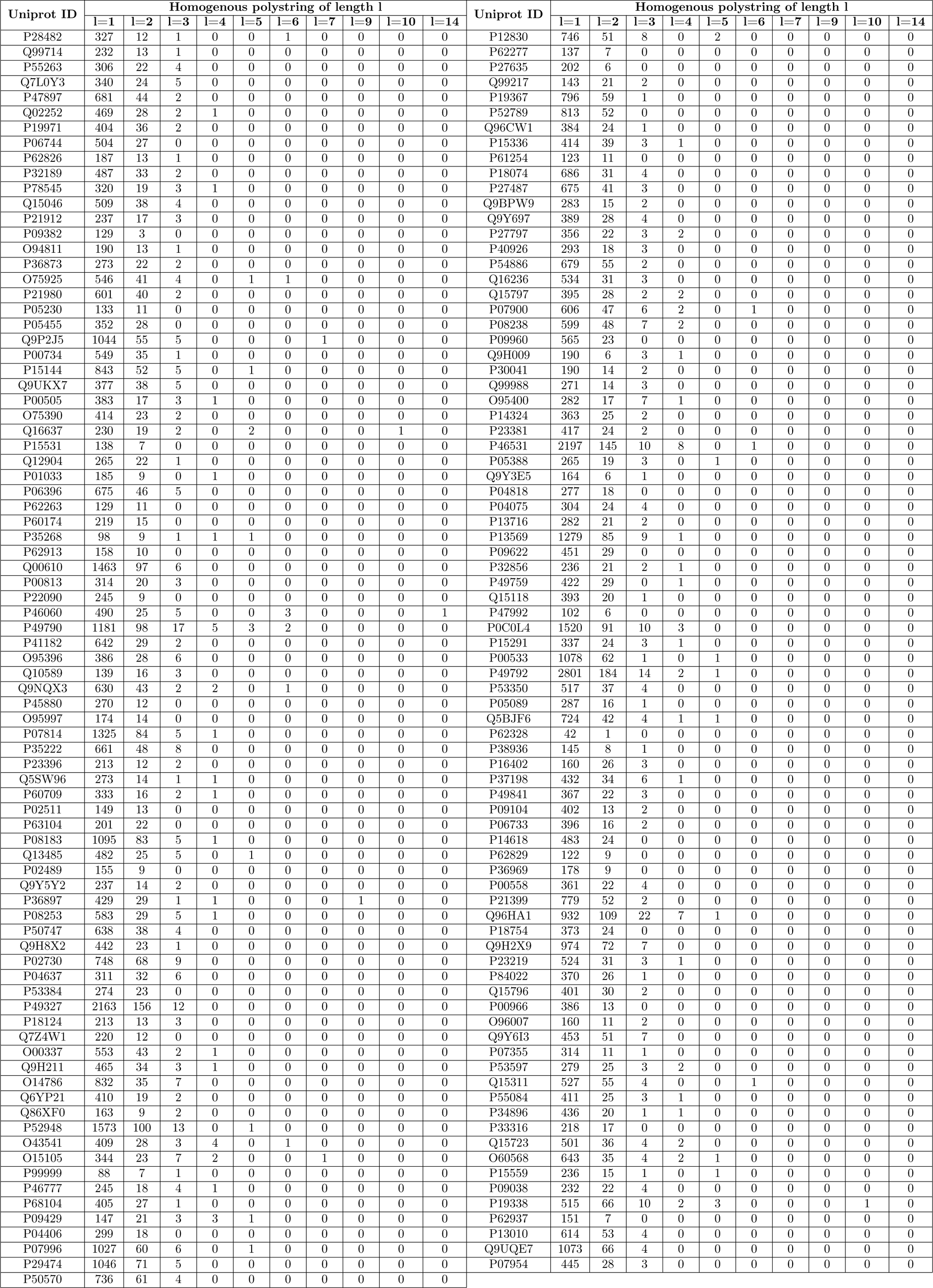
Frequency of homogeneous poly-string of length 1, 2, … 14 doe each moonlighting proteins.

Additionally, poly-strings length of 10 appeared in sequences Q16637 and P19338, consisting of prolin and glutamic acid, respectively, each with a frequency of 1. Furthermore, P36897 contained a poly-string length of 9 comprised of alanine. Notably, sequences Q9P2J5 and O15105 exhibited poly-strings length of 7, featuring glutamic acid and glycine, respectively, both occurring once. Poly-strings lengths of 3, 4, 5, and 6 were identified in 130, 43, 18, and 9 many number of sequences, respectively. For a single sequence, poly-strings length of 7, 9, 10, and 14 were noticed with the highest frequency 1, while poly-string length of 3, 4, 5, 6 were noticed with the highest frequency 22, 8, 3, and 3, respectively. Noteworthy observations included three poly-strings of length 6 in P46060, all composed of glutamic acid, and three poly-strings of length 5 in both P49790 and P19338, originating from distinct amino acids, as detailed in **Supplementary file-2**. Furthermore, it was revealed that all 165 moonlighting proteins contained homogeneous poly-strings of lengths 1 and 2. Impressively, these were most prevalent in P49792, with frequencies of 2801 and 184, respectively. This observation emphasizes the common occurrence of short poly-strings composed of individual amino acids or consecutive pairs within these proteins.

The diversity observed in poly-string lengths for various amino acids underscores the intricate arrangement and variability of amino acids within the moonlighting proteins, highlighting the complexity of their structural composition.

### 4.3. Polar, non-polar residue profiles of moonlighting proteins

The percentage distributions of polar and non-polar residues were computed for each moonlighting protein, as presented in Figure 5. The analysis revealed that the average ratio of polar to non-polar residues was 1.0061*±*0.2462. Notably, five sequences—Q9Y5Y2, Q99217, O00337, Q99714, and P19971—exhibited relatively lower ratios of 0.632, 0.63, 0.606, 0.601, and 0.58, respectively. On the higher end, the sequences Q9UQE7, Q5BJF6, P49759, P61254, P09429, and P62328 displayed elevated ratios of polar to non-polar residues, measuring 1.57, 1.62, 1.65, 1.79, 1.83, and 2.384, respectively. Furthermore, a distinct observation was made regarding the sequences P28482, P55263, P06744, and Q6YP21, where the percentages of polar and non-polar residues were exactly identical, resulting in a ratio of 1 for each sequence, as depicted in Figure 5. This uniformity in the ratios underscores a specific compositional characteristic shared among these sequences.

**Figure 5:**
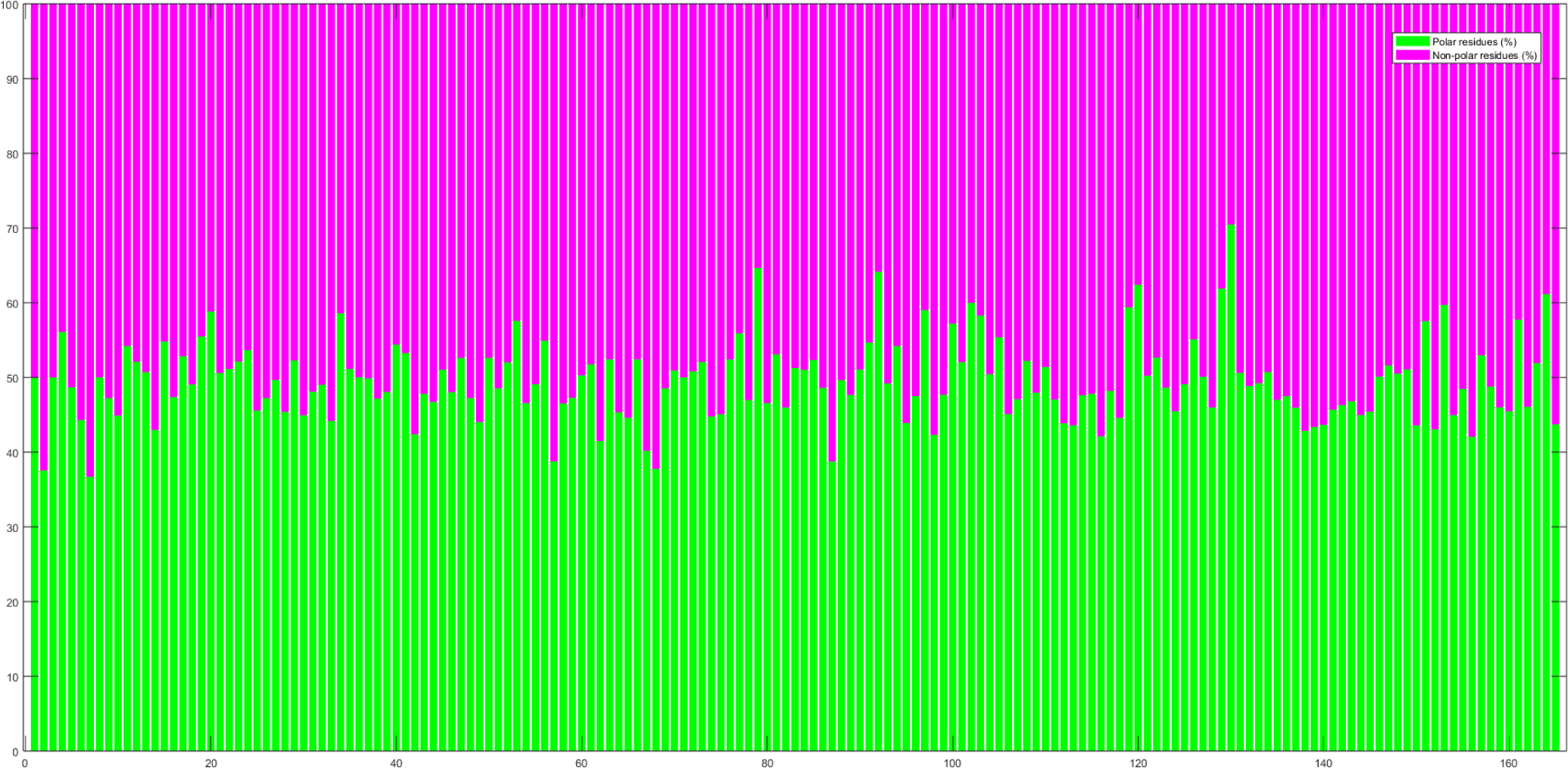
Percentages of polar, non-polar residues in moonlighting proteins

The analysis delineates the distribution of polar and non-polar residues in moonlighting proteins, revealing a range of ratios that vary across different sequences. The observed variations provide insights into the potential functional and structural diversity inherent in these proteins.

#### 4.3.1. Change response sequences of polar, non-polar profiles

Figure 6 shows the relative frequency distribution of four changes namely PN, NP, PP, and NN. Percentages of polar to polar residue changes for the sequences P61254, P09429, P49759, and P62328 were significantly high compared to others namely 40.28%, 42.06%, 42.65%, and 51.16%, respectively. Intriguingly, among these, P62328 displayed the lowest percentages of ‘PN’ (18.6%), ‘NN’ (9.3%), and ‘NP’ (20.9%) changes. This emphasizes P62328’s distinct residue change profile and suggests potential sequence-specific functional or structural implications. The observed variations in polar residue substitutions underscore the sequence-specific nature of residue changes and hint at their potential functional significance in these proteins.

**Figure 6:**
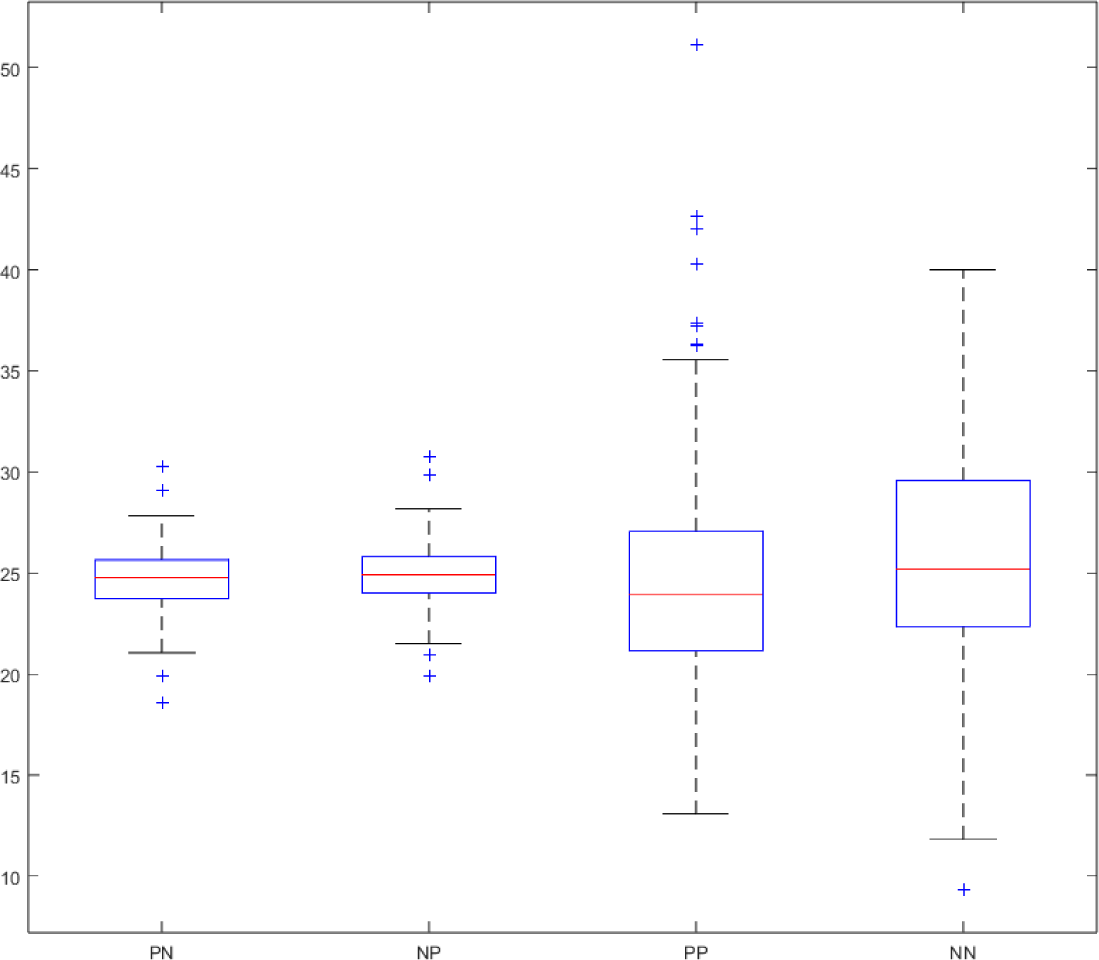
Box-plot of the relative frequency of PN, NP, PP, and NN changes in moonlighting proteins

Based on a distance threshold of 5, an analysis of 160 moonlighting sequences led to the formation of eleven distinct clusters, as depicted in the dendrogram (Figure 7 (A)) and summarized in Table 7. The most substantial cluster was comprised of 50 sequences, highlighted by a brown rectangular box in Figure 7 (A). Notably, sequence P62328 exhibited considerable dissimilarity from the majority of sequences. Particularly, P62328 displayed a distance greater than 28 from 131 other moonlighting sequences.

**Figure 7:**
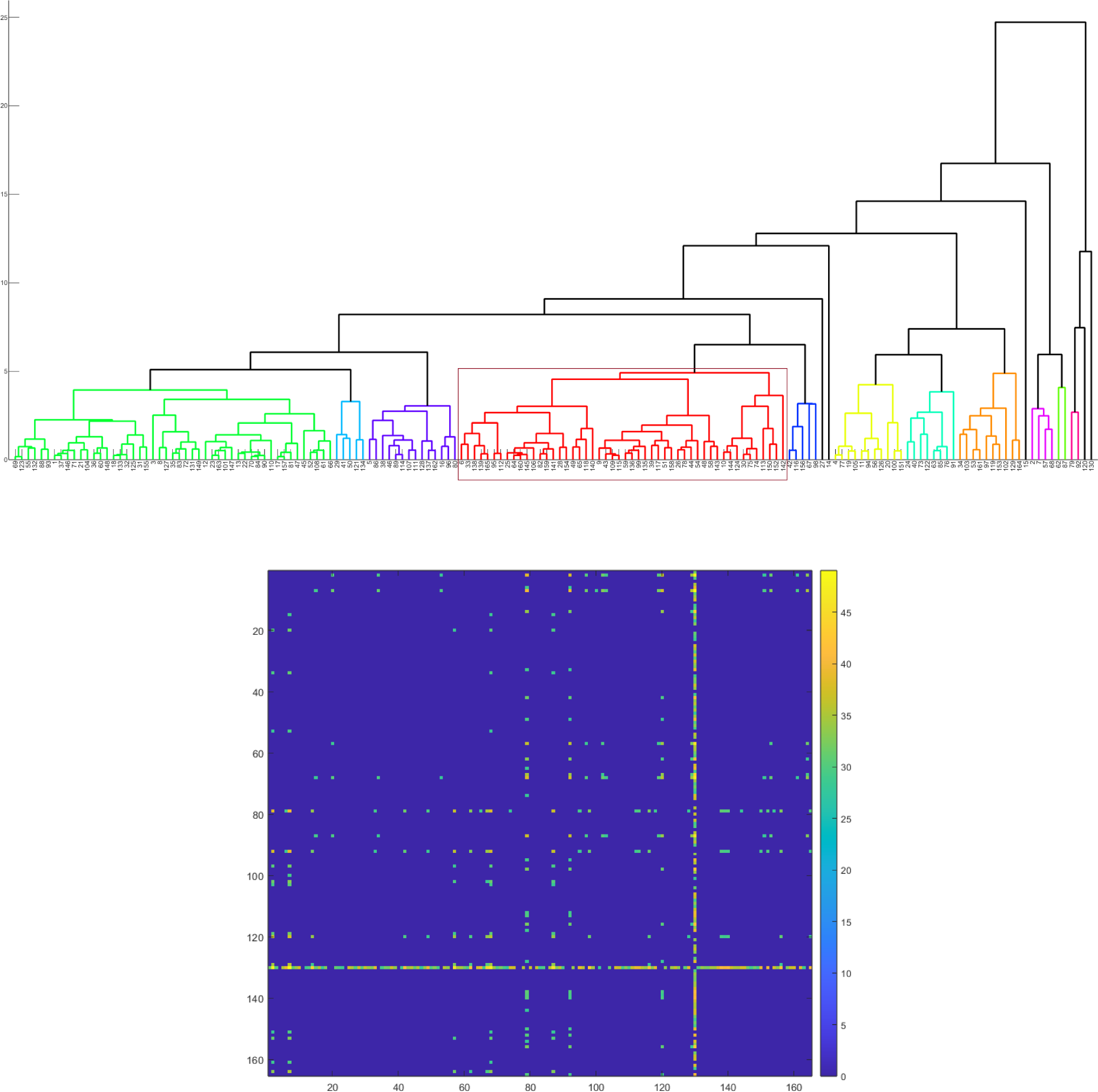
(A): Phylogenetic relationship among the moonlighting proteins based on relative frequency of PP, NP, PP, and NN changes as obtained from polar, non-polar profiles. (B): Distance matrix consisting distances between pair of relative frequency (of PP, NP, PP, and NN changes) vectors corresponding to each moonlighting protein (distance less than 28 considered to be zero).

**Table 7:**
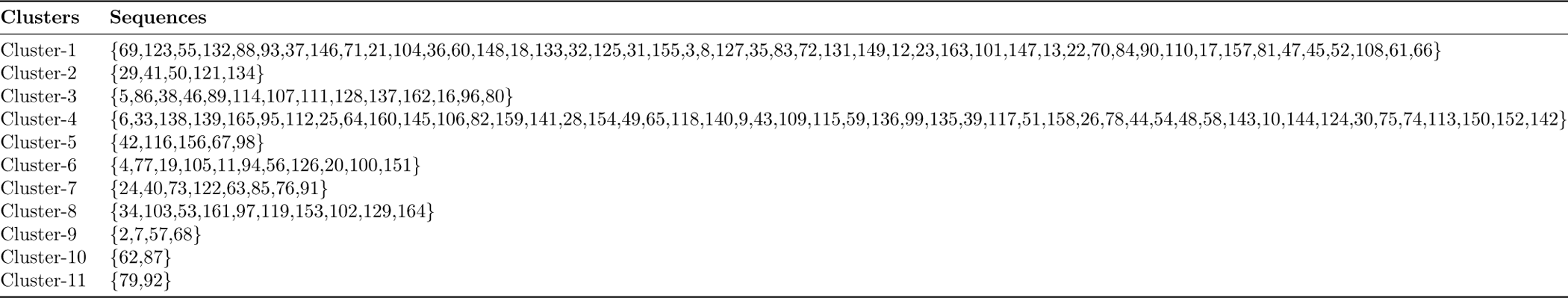
Clusters of moonlighting proteins based on relative frequency of PP, NP, PP, and NN changes as obtained from Polar, non-polar profiles.

| Clusters | Sequences |
| --- | --- |
| Cluster-1 | {69,123,55,132,88,93,37,146,71,21,104,36,60,148,18,133,32,125,31,155,3,8,127,35,83,72,131,149,12,23,163,101,147,13,22,70,84,90,110,17,157,81,47,45,52,108,61,66} |
| Cluster-2 | {29,41,50,121,134} |
| Cluster-3 | {5,86,38,46,89,114,107,111,128,137,162,16,96,80} |
| Cluster-4 | {6,33,138,139,165,95,112,25,64,160,145,106,82,159,141,28,154,49,65,118,140,9,43,109,115,59,136,99,135,39,117,51,158,26,78,44,54,48,58,143,10,144,124,30,75,74,113,150,152,142} |
| Cluster-5 | {42,116,156,67,98} |
| Cluster-6 | {4,77,19,105,11,94,56,126,20,100,151} |
| Cluster-7 | {24,40,73,122,63,85,76,91} |
| Cluster-8 | {34,103,53,161,97,119,153,102,129,164} |
| Cluster-9 | {2,7,57,68} |
| Cluster-10 | {62,87} |
| Cluster-11 | {79,92} |

In the context of pairwise distances, five proteins—Q99714, P19971, Q9Y5Y2, O00337, and Q99217—were identified as the furthest from P62328. The calculated distances ranged from 45 to 49, with 49 representing the highest pairwise distance among all comparisons. These insights were visualized through a distance matrix, presented in Figure 7 (B), with a defined threshold cut off at 28.

These findings underscore the diversity and relationships among the analyzed moonlighting sequences, revealing distinct clusters and the unique positioning of P62328 in relation to other sequences. The pairwise distances further highlight specific sequences that stand out in terms of dissimilarity from P62328, contributing to a comprehensive understanding of the sequence relationships within this dataset.

### 4.4. Acidic, Basic, Neutral residue based phylogenetic relationship

Based on the data illustrated in Figure 8, the percentages of acidic, basic, and neutral residues were calculated for each moonlighting protein. In this context, the analysis encompassed 165 sequences, exhibiting varying percentages of neutral residues spanning from 50.2% to 93.2%. The calculated ratio of acidic to basic residue percentages was determined to be 1.03*±*0.29, indicating a relatively balanced distribution between acidic and basic residues across the dataset. Furthermore, it was observed that the ratio of acidic to basic residues precisely equaled 1 for three specific sequences: P62826, P06733, and P18754.

**Figure 8:**
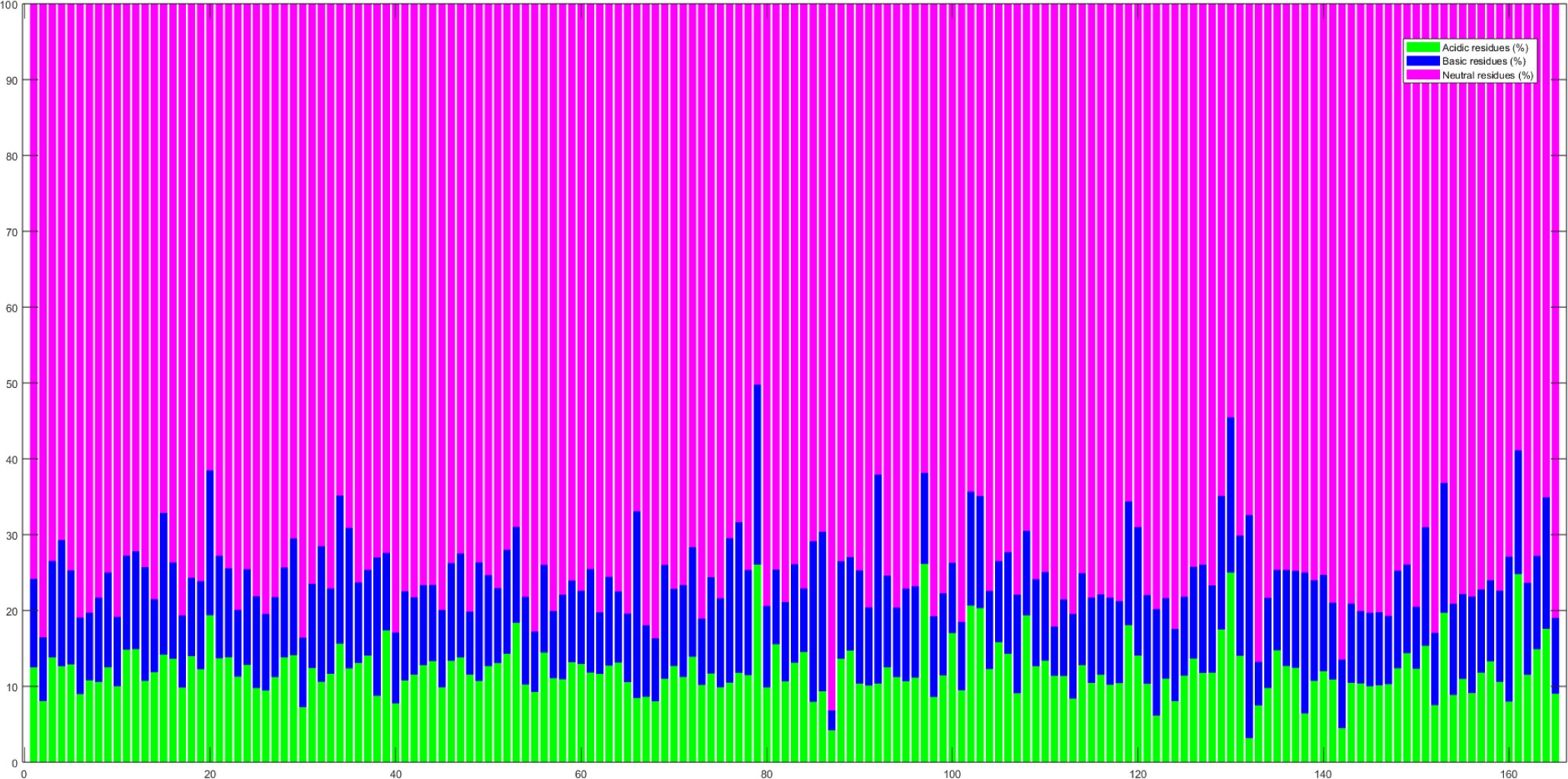
Percentages of acidic, basic, neutral residues in moonlighting proteins

The sequence P16402 stood out with distinct residue composition characteristics. It possessed the lowest percentage of acidic residues, measuring 3.17%, and concurrently displayed the highest percentage of basic residues, at 29.41%.

These findings collectively emphasize the diversity in residue compositions across the moonlighting proteins. The ratio of acidic and basic residues, as well as the particular characteristics of sequences such as P16402, provide insights into the potential functional implications of specific residue types within these proteins.

#### 4.4.1. Change response sequences of acidic, basic, and neutral profiles

The relative frequency distribution of nine different residue changes, denoted as BA, NA, AA, BB, NB, AB, BN, NN, and AN, was visualized in Figure 9. Among all the sequences, the highest percentages were observed for the BA (AB) changes, amounting to 4.74% (11.63%) in Q15311 (P62328). Notably, P62328 exhibited the highest percentages of NA (AN) changes, standing at 23.25% (13.95%), while P16402 had the lowest percentages, at 2.72% (1.81%) respectively.

**Figure 9:**
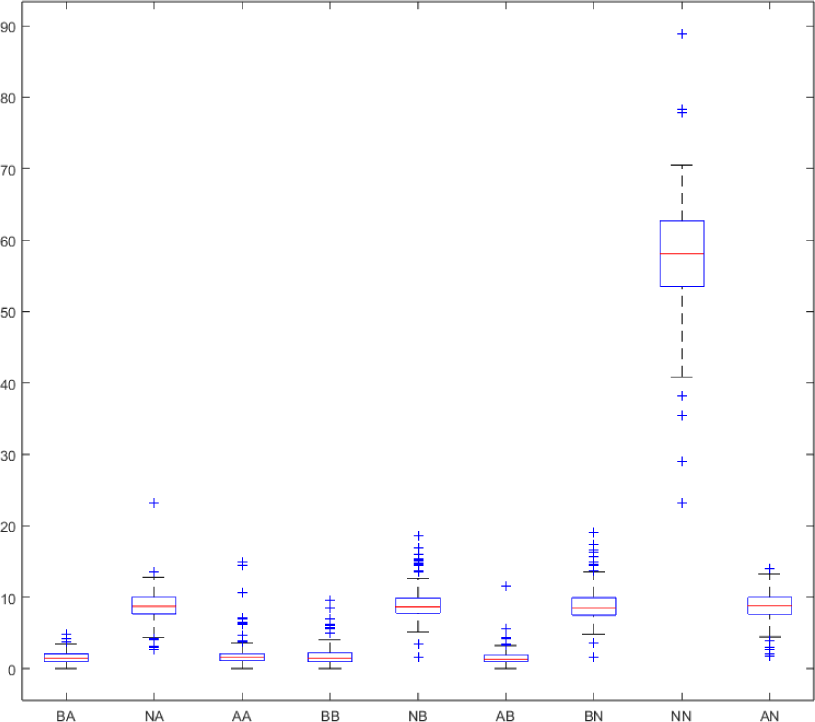
Box-plot of the relative frequency of all nine changes in moonlighting proteins

In terms of specific changes, P09429 displayed the highest percentage of AA changes, accounting for 14.95% of its residues. P16402, on the other hand, exhibited the highest percentage of BB changes, comprising 9.54% of its residues. Interestingly, P16402 also stood out with the highest percentages of NB (BN) changes, reaching 18.64% (19.1%), whereas Q99217 had the lowest percentages, both recorded at 1.58% (1.58%). The highest percentage of NN changes was found in Q99217, constituting 88.95% of its residues. Conversely, P62328 showcased the lowest percentage of NN changes, at 23.25%.

These observations provide insights into the distinct residue change patterns across various sequences. The highest and lowest percentages of specific changes highlight the variability in these sequences and potentially suggest functional or structural significance associated with certain types of residue alterations.

Following change responses were absent in case of following moonlighting sequences:

- Basic to Acidic: Q9Y3E5
- Acidic to Acidic: P16402, P47992, Q7Z4W1, P09382, and P62328
- Basic to Basic: P09382
- Acidic to Basic: P09382, Q99988, and P62829

Under a distance threshold of 5, the analysis led to the formation of twelve distinct clusters comprising 144 moonlighting sequences. This information is visually depicted in the dendrogram (Figure 10 (A)) and summarized in Table 8. Notably, the largest cluster contained 56 sequences, indicated by a brown rectangular box in Figure 10 (A). Several sequences demonstrated considerable distance from others, reflecting their distinct positions within the dataset. Specifically, sequences P05455, P09429, Q99217, P62328, and P19338 were situated more than 30 distances away from 37, 117, 115, 150, and 32 sequences, respectively. The maximum calculated distance of 72.27 was observed between sequences P62328 and Q99217, underscoring the significant dissimilarity between these two.

**Figure 10:**
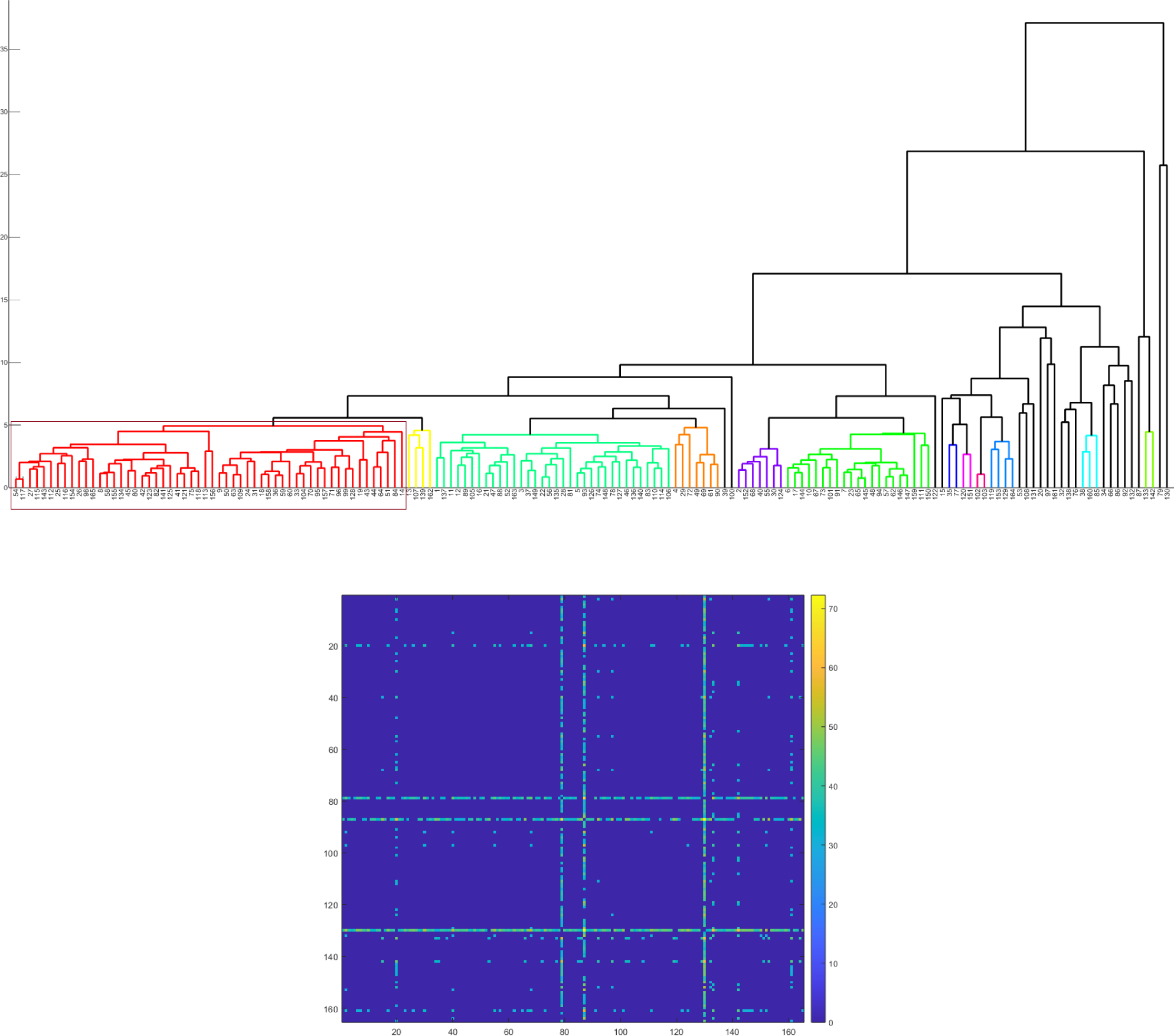
(A): Phylogenetic relationship among the moonlighting proteins based on relative frequency of BA, NA, AA, BB, NB, AB, BN, NN, and AN changes as obtained from acidic, basic, and neutral profiles. (B): Distance matrix consisting distances between pair of relative frequency (of BA, NA, AA, BB, NB, AB, BN, NN, and AN changes) vectors corresponding to each moonlighting protein (distance less than 30 considered to be zero).

**Table 8:** Clusters of moonlighting proteins based on relative frequency of BA, NA, AA, BB, NB, AB, BN, NN, and AN changes as obtained from acidic, basic, and neutral profiles.

| Cluster | Sequences |
| --- | --- |
| Cluster-1 | {54,117,27,115,143,112,25,116,154,26,98,165,8,58,155,134,45,80,42,123,82,141,125,41,121,75,118,113,156,9,50,63,109,24,31,18,158,36,59,60,33,104,70,95,157,71,96,99,128,19,43,44,64,51,84,14} |
| Cluster-2 | {13,107,139,162} |
| Cluster-3 | {1,137,11,12,89,105,16,21,47,88,52,163,3,37,149,22,56,135,28,81,5,93,126,74,148,78,127,46,136,140,83,110,114,106} |
| Cluster-4 | {4,29,72,49,69,61,90} |
| Cluster-5 | {2,152,68,40,55,30,124} |
| Cluster-6 | {6,17,144,10,67,73,101,91,7,23,65,145,48,94,57,62,146,147,159,111,150} |
| Cluster-7 | {35,77} |
| Cluster-8 | {120,151} |
| Cluster-9 | {102,103} |
| Cluster-10 | {119,153,129,164} |
| Cluster-11 | {38,160,85} |
| Cluster-12 | {133,142} |

These findings were effectively captured through a distance matrix, illustrated in Figure 10 (B), with a designated threshold cutoff set at 30.

Collectively, these results provide valuable insights into the structural relationships and variations among the analyzed moonlighting sequences. The identified clusters and pairwise distances contribute to a comprehensive understanding of the sequence associations within the dataset.

### 4.5. Intrinsic protein disorder analysis of moonlighting proteins

The intrinsic protein disorder probability of each residue was calculated for every moonlighting sequence. The percentages of disordered residues, highly flexible residues, moderately flexible residues, and other residues in each moonlighting protein were determined and presented in Figure 11. It’s worth noting that one sequence, P36969, was excluded from the intrinsic protein disorder analysis due to the presence of an ambiguous character ‘U’.

**Figure 11:**
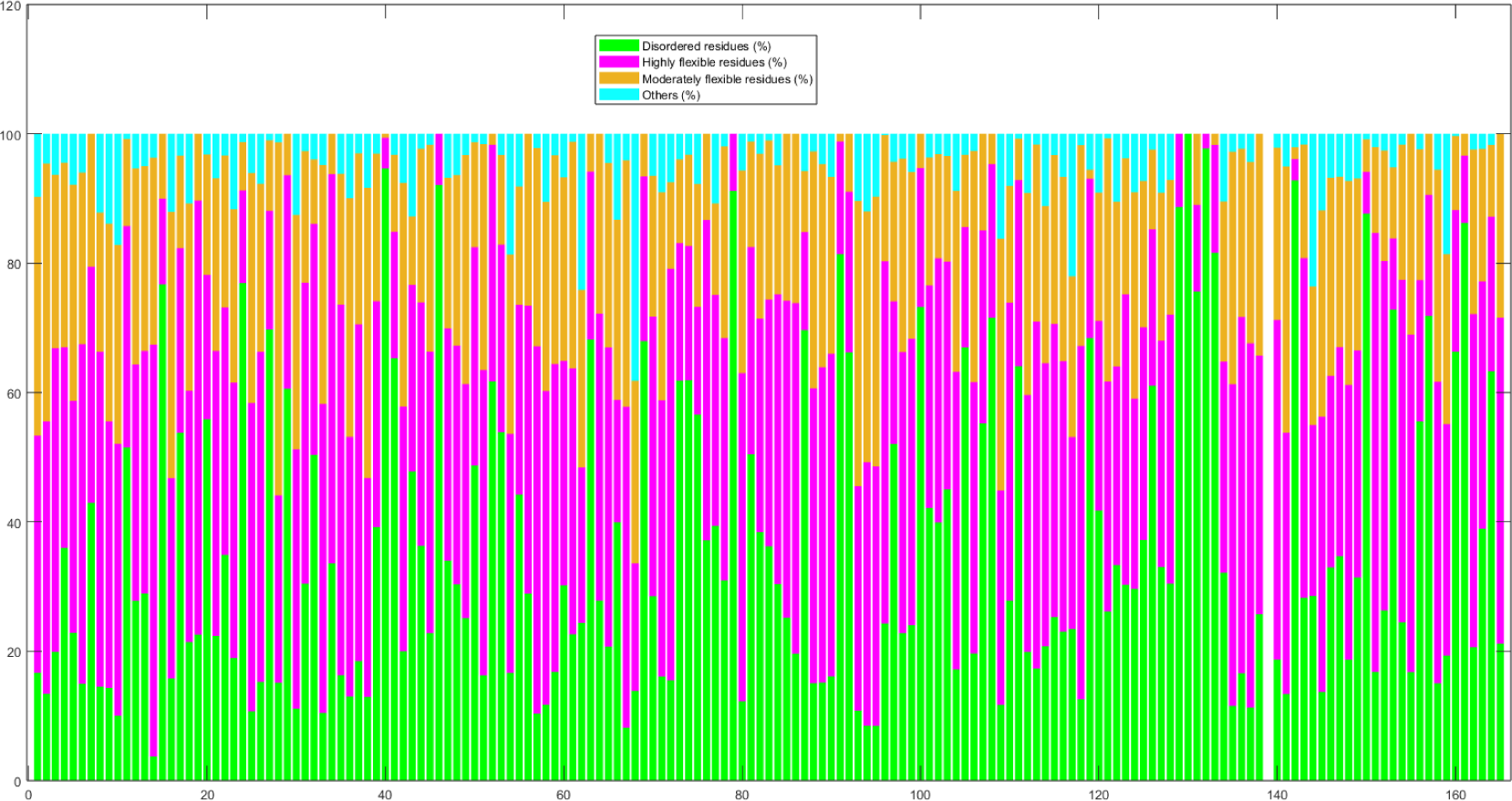
Percentages of disordered, highly flexible, moderately flexible and other residues in moonlighting proteins

An interesting observation was made regarding the moonlighting sequence P62328, which was found to consist entirely of disordered residues. Additionally, six moonlighting proteins exhibited a notably high percentage of disordered residues, namely P16402 (97.74%), P49790 (94.64%), Q96HA1 (92.79%), O95997 (92.08%), P09429 (91.16%), and Q5BJF6

(88.66%). Conversely, these proteins exhibited a significantly low percentage of highly flexible residues, with P16402 (2.26%), Q96HA1 (3.28%), P49790 (4.75%), Q9Y6I3 (6.42%), O95997 (7.92%), P09429 (8.84%), P19338 (10.42%), and Q15311 (10.99%).

In contrast, four sequences, namely Q86XF0 (63.64%), P09382 (63.70%), P05230 (67.10%), and P07355 (67.85%), displayed a notably high percentage of highly flexible residues. Remarkably, five moonlighting sequences, O95997, P09429, Q5BJF6, P62328, and P16402, did not exhibit any moderately flexible residues. The highest and lowest (non-zero) percentages of moderately flexible residues were observed in P15531 (54.61%) and P49790 (0.61%), respectively.

#### 4.5.1. Global disorder status of human moonlighting proteins

Global disorder analysis revealed that human moonlighting proteins contain noticeable levels of intrinsic disorder. This is illustrated by Figure 12A representing the results of the classification of the disorder status of these proteins based on the outputs of the per-residue disorder predictor PONDR® VSL2 in a form of the MDS vs. PPIDR dependence. This plot shows distribution of the moonlighting proteins within segments containing mostly ordered (blue and light blue), moderately disordered (pink and light pink), or mostly disordered (red) proteins. Based on these classifications, none of the moonlighting proteins was predicted as ordered by both MDS and PPIDR (dark blue segment is empty), and only 4 (2.4%) of these were predicted as mostly ordered based on their MDS values (light blue segment). Therefore, remaining moonlighting proteins are expected to be either moderately or highly disordered. In fact, Figure 12A shows that 50.3% of these proteins were predicted as moderately disordered and containing noticeable levels of flexible or disordered residues/regions based on both their MDS and PPIDR values (dark pink segment). Additional 21.2% proteins were expected to be moderately disordered based on their MDS values (light pink segment), whereas 26.1% of the human moonlighting proteins are expected to be highly disordered (red segment). Importantly, most of these proteins (42 of 43) have PPIDR ≥ 50% and MDS ≥ 0.5.

**Figure 12:**
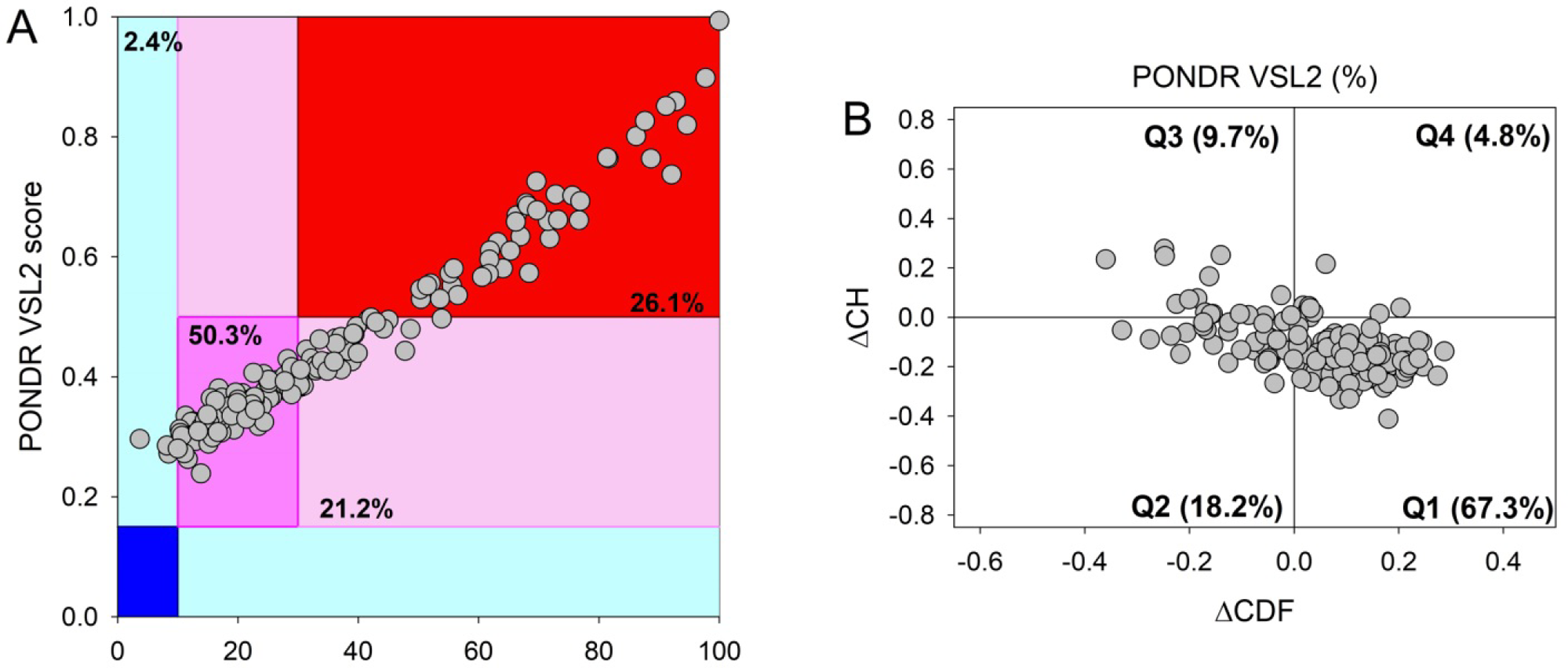
Global disorder analysis of human moonlighting proteins. A. PONDR® VSL2 output for 165 human moonlighting proteins. PONDR® VSL2 score is the mean disorder score (MDS) for a protein, which is calculated as a sequence-length normalized sum of its perresidue scores. PONDR VSL2 (%) is a percent of predicted intrinsically disordered residues (PPIDR); i.e., percent of residues with disorder scores above 0.5. Color blocks indicate regions in which proteins are mostly ordered (blue and light blue), moderately disordered (pink and light pink), or mostly disordered (red). If the two parameters agree, the corresponding part of the background is dark (blue or pink), whereas light blue and light pink reflect areas in which only one of these criteria applies. The boundaries of the colored regions represent arbitrary and accepted cutoffs for MDS (y-axis) and the percentage of predicted disordered residues (PPDR; x-axis). Of the 165 human moonlighting proteins, 44 (26.6%) are predicted to be highly disordered, 118 (71.5%) demonstrate moderate disorder or conformational flexibility, and only 4 proteins (2.4%) are predicted to be highly ordered. B. Charge-hydropathy and cumulative distribution function (CH-CDF) plot for 165 human moonlighting proteins. The Y-coordinate is calculated as the distance of the corresponding protein from the boundary in the CH plot. The X-coordinate is calculated as the average distance of the corresponding protein’s CDF curve from the CDF boundary. The quadrant where the proteins are located determines their classification. Q1, 111 proteins (67.3%) predicted to be ordered by CDF and compact/ordered by CH-plot. Q2, 30 proteins (18.2%) predicted to be ordered/compact by CH-plot and disordered by CDF. Q3, 16 proteins (9.7%) predicted to be disordered by CH-plot and CDF. Q4, 8 proteins (4.8%) predicted to be disordered by CH-plot and ordered by CDF

Further support for relatively high disorder status of human moonlighting proteins was provided by combining the outputs of two binary predictors charge-hydropathy (CH) plot and cumulative distribution function (CDF) analysis that classify proteins as mostly ordered or mostly disordered (see Figure 12B). The Result of the CH-CDF plot provides useful means for more detailed characterization of the global disorder status of proteins, classifying them as mostly ordered, molten globule-like or hybrid, or highly disordered (see Materials and Methods section). Figure 12B shows that 67.3% of the moonlighting proteins are located within the quadrant Q1 (bottom right corner) containing proteins predicted to be ordered by both predictors, whereas 32.7% of these proteins are located outside the quadrant Q1 and can be considered as proteins with high disorder levels. Figure 12B also shows that the quadrant Q2 (bottom left corner) corresponding to molten globular or hybrid proteins predicted to be ordered/compact by the CH-plot and disordered by the CDF analysis includes 18.2% of these proteins. On the other hand, 9.7% of moonlighting proteins, being located in the quadrant Q3 (top left corner), are predicted as highly disordered by both predictors, whereas the quadrant Q4 (top right corner) contains 4.8% proteins predicted as disordered by CH-plot and ordered by CDF analysis.

#### 4.5.2. Change response sequences of disordered, highly flexible, moderately flexible, and other residues profiles

In Figure 13, the relative frequency distribution of sixteen changes derived from the change response sequence based on intrinsic disorder profiles is presented.

**Figure 13:**
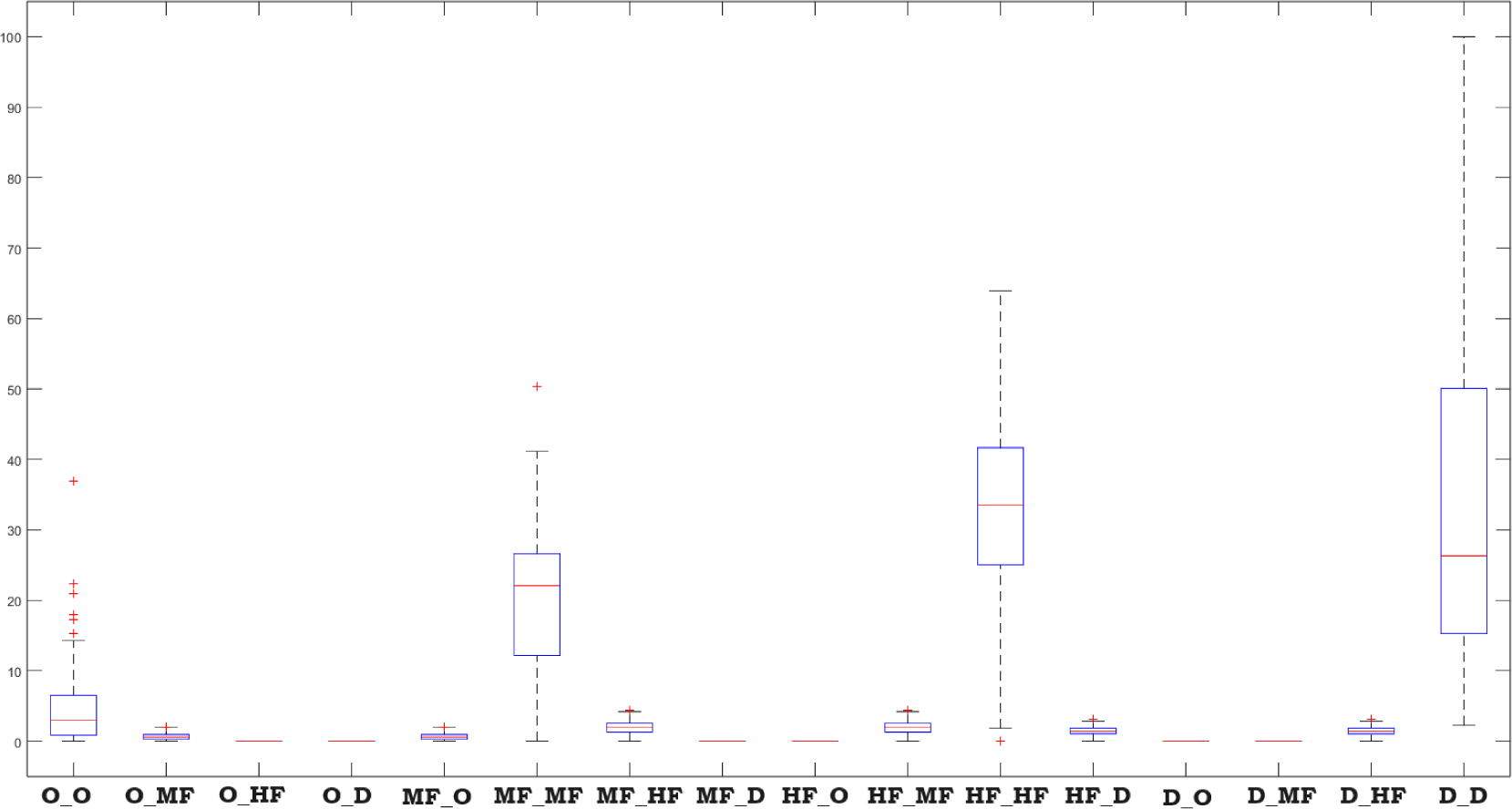
Box-plot of the percentages of disordered, highly flexible, moderately flexible and other residues changes in moonlighting protein sequences

Notably, certain moonlighting protein sequences, specifically O00337, P02730, Q9H2X9, and P13569, exhibited significantly high percentages of O_O changes of residues, with values of 36.88%, 22.42%, 22.41%, and 20.90%, respectively. This suggests a notable propensity for these sequences to transition between other states.

An intriguing observation is that none of the 164 moonlighting protein sequences displayed changes from disordered to moderately flexible, disordered to other, highly flexible to other, moderately flexible to disordered, other to disordered, or other to highly flexible residues. This lack of transitions between these specific states is worth noting. Furthermore, it was noticed that the highest amount (50.33%) of changes from moderately flexible to moderately flexible (self-transition) was noticed in P15531 (Figure 13). P07355 shows the highest percentage of (63.91%) of changes from highly flexible to highly flexible residues. Figure 13 provides valuable insights into the dynamic changes in intrinsic disorder profiles among the moonlighting protein sequences, highlighting specific sequences with distinctive disorder transition patterns.

An analysis of 159 moonlighting sequences, using a distance threshold of 15, resulted in the formation of eleven distinct clusters, which can be observed in the dendrogram (Figure 14 (A)) and are summarized in Table 9. The most substantial cluster, Cluster-6, consisted of 53 sequences and is highlighted by a brown rectangular box in Figure 14 (A).

**Figure 14:**
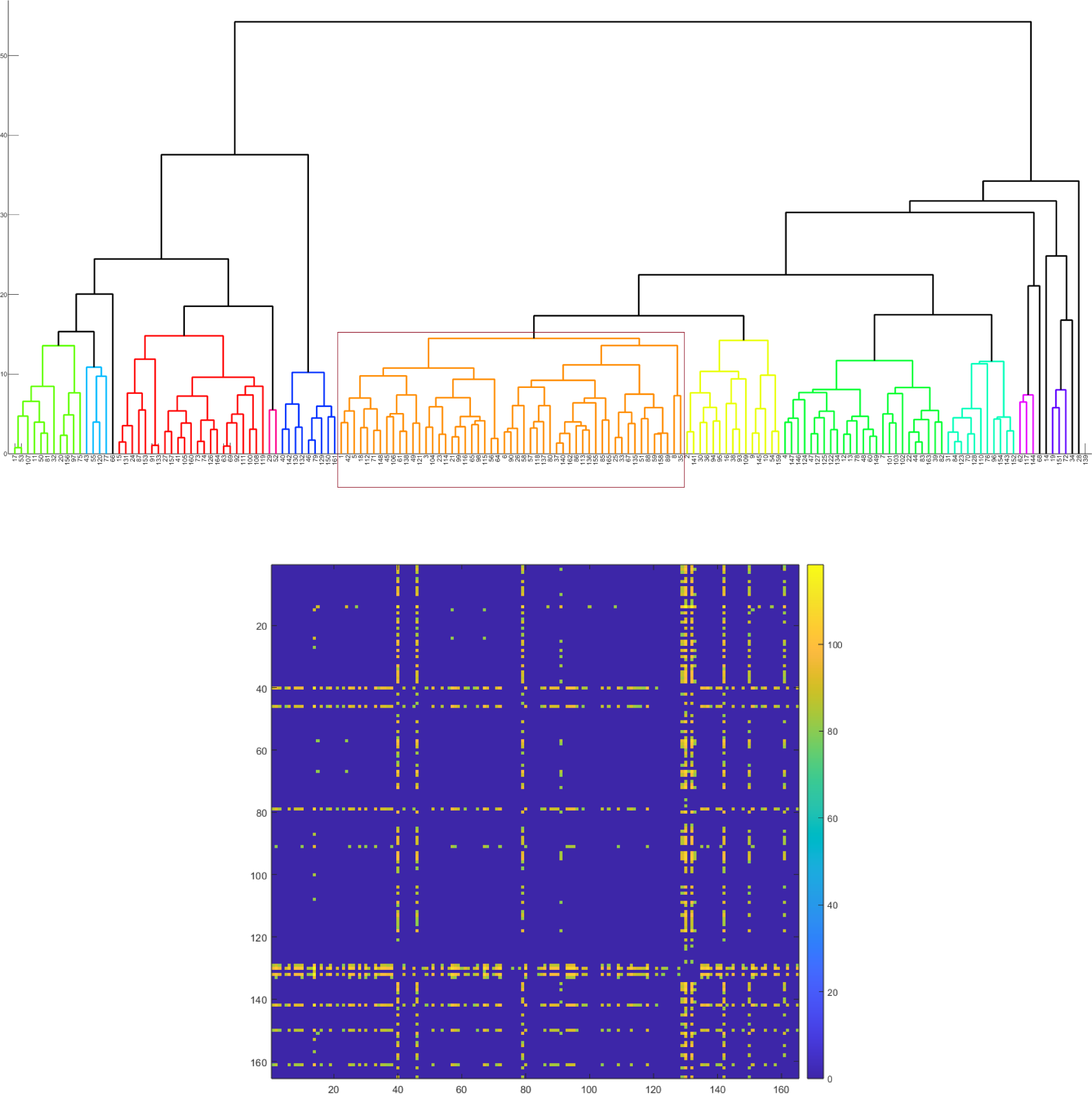
(A): Phylogenetic relationship among the moonlighting proteins based on percentages of disordered, highly flexible, moderately flexible and other residues. (B): Distance matrix consisting distances between pair of percentages of disordered, highly flexible, moderately flexible and other residues corresponding to each moonlighting protein (distance less than 80 considered to be zero).

**Table 9:**
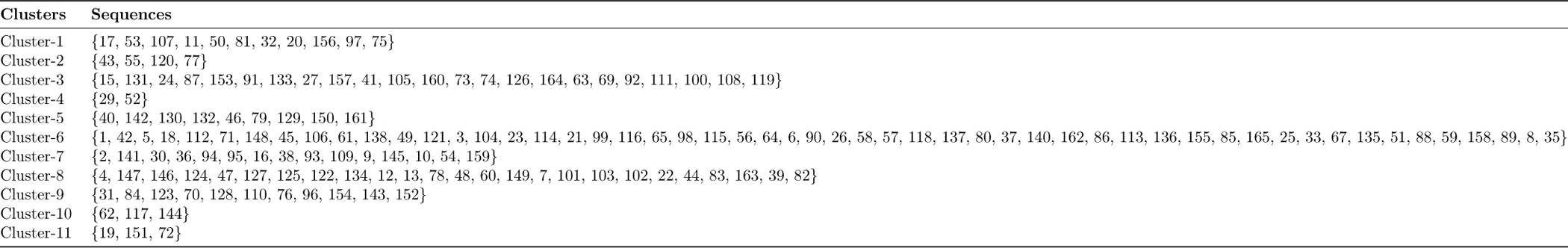
Clusters of moonlighting proteins based on the change response profiles of disordered, highly flexible, moderately flexible, and other residues in moonlighting proteins.

Interestingly, certain sequences, such as P49790, O95997, P09429, P62328, P16402, Q96HA1, Q9Y6I3, and P19338, were notably distant from a considerable number of other sequences, with distances greater than 80. Specifically, they were distant from 83, 73, 68, 98, 90, 80, 67, and 49 sequences, respectively. Among all possible pairwise distances, the maximum distance recorded was 118.34 for the pair P09382 and P62328. These insights were visually represented in a distance matrix, as shown in Figure 7 (B), with a defined threshold cutoff set at 80.

### 4.6. Phylogenetic relationship based on structural and physicochemical features

Under a distance threshold of 6.5, a comprehensive analysis revealed the formation of eleven distinct clusters encompassing 62 moonlighting sequences. This partitioning of sequences is visually represented in the dendrogram (Figure 15 (A)) and concisely detailed in Table 10. Notably, the most substantial cluster comprised 33 sequences, which is highlighted by a brown rectangular box in Figure 15 (A).

**Figure 15:**
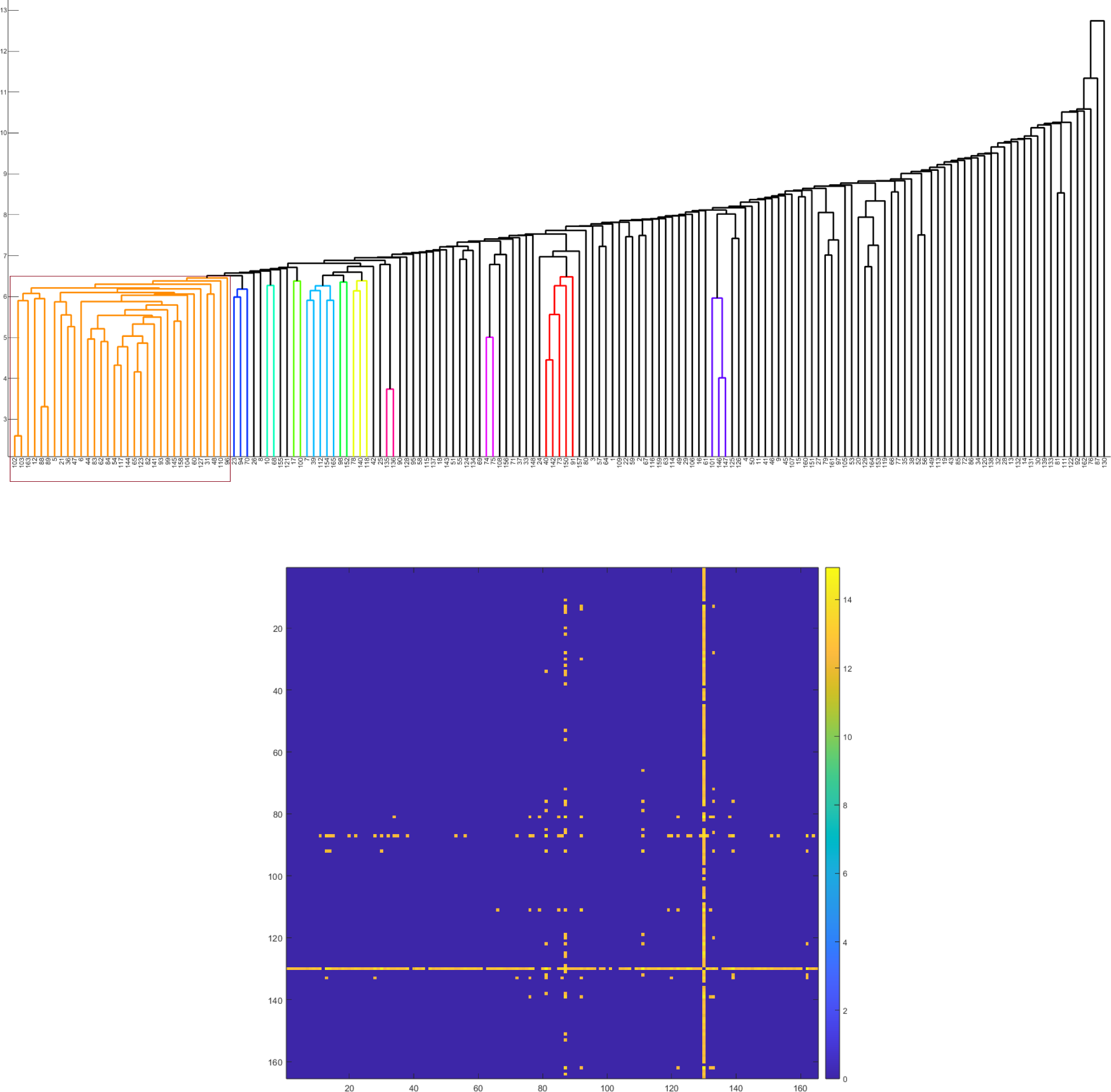
(A): Phylogenetic relationship among the moonlighting proteins based on structural and physicochemical features. (B): Distance matrix consisting distances between pair of structural and physicochemical quantitative vectors corresponding to each moonlighting protein (distance less than 12 considered to be zero).

**Table 10:** Clusters of moonlighting proteins based on on structural and physicochemical features.

| Clusters | Sequences |
| --- | --- |
| Cluster-1 | {102,103,163,12,88,89,5,21,36,47,6,44,83,62,84,54,117,144,65,123, 82,141,93,99,145,158,104,60,127,31,48,110,96} |
| Cluster-2 | {23,94,70} |
| Cluster-3 | {10,68} |
| Cluster-4 | {17,100} |
| Cluster-5 | {7,39,112,154,165} |
| Cluster-6 | {98,152} |
| Cluster-7 | {78,140,118} |
| Cluster-8 | {135,136} |
| Cluster-9 | {74,75} |
| Cluster-10 | {40,142,73,150,91} |
| Cluster-11 | {101,146,147} |

The analysis further revealed intriguing patterns of sequence distances. Specifically, sequences Q99217 and P62328 exhibited considerable dissimilarity from other sequences, as they were separated by a distance of more than 12, from 36 and 149 sequences, respectively. The maximum observed distance of 14.94 was recorded between sequences P07996 and P62328, underscoring a notable dissimilarity between these two sequences. These insights were concisely captured in a distance matrix, as illustrated in Figure 15 (B), with a predefined cutoff threshold of 12.

Collectively, the results shed light on the structural relationships and distinctions among the evaluated moonlighting sequences. The formation of clusters and the identification of specific sequences with notable dissimilarities contribute to an enhanced understanding of sequence associations within the dataset.

### 4.7. Cumulative clustering of moonlighting proteins

Utilizing a distance threshold of 150, an analysis of 156 moonlighting sequences resulted in the formation of six distinct clusters, as illustrated in the dendrogram (see Figure 16(A)) and summarized in Table 11. The most extensive cluster, Cluster-3, encompassed 98 sequences and is highlighted within a brown rectangular box in Figure 16(A).

**Figure 16:**
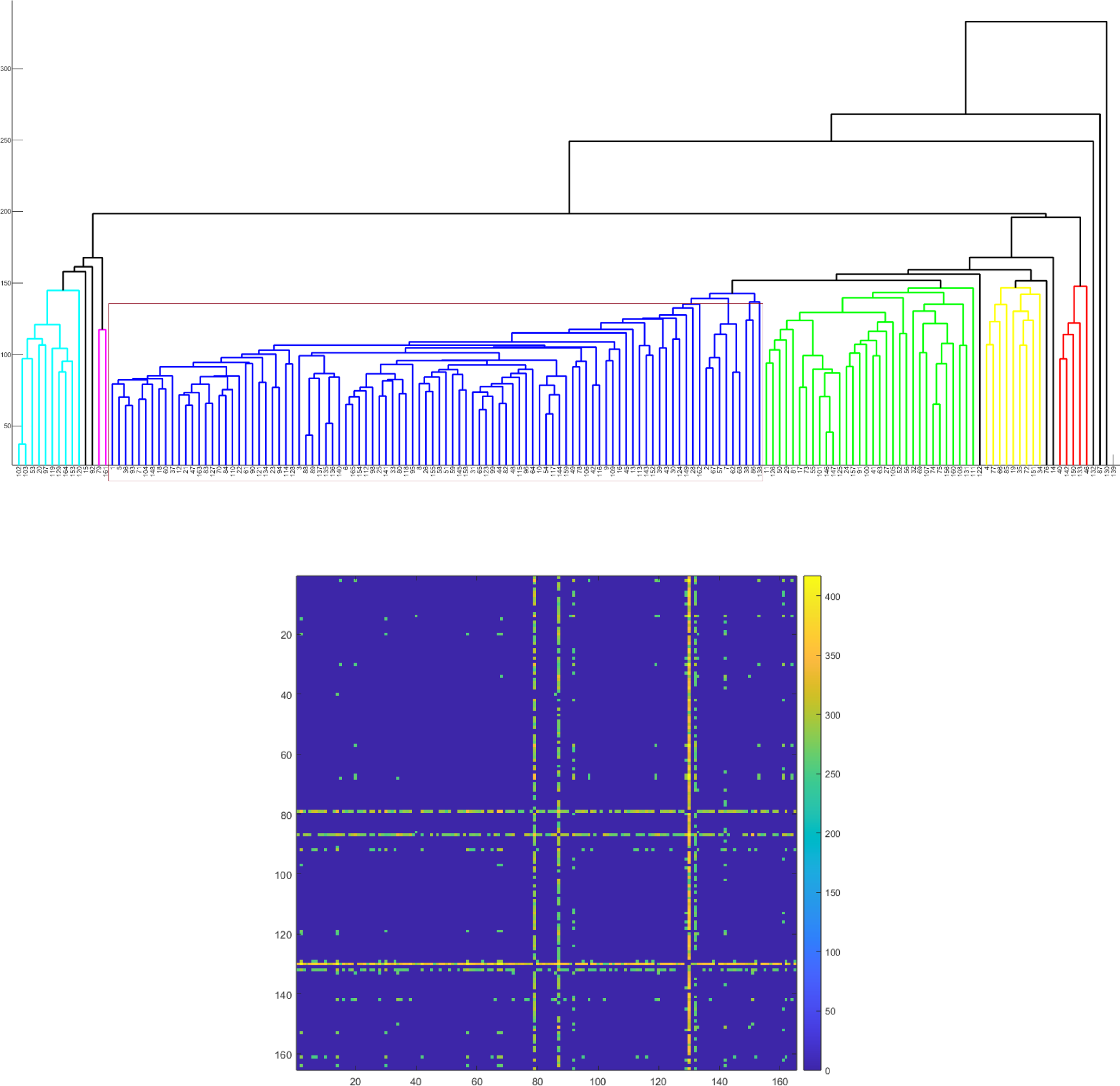
(A): Phylogenetic relationship among the moonlighting proteins from cumulative clustering. (B): Corresponding distance matrix (distance less than 250 considered to be zero).

**Table 11:** Cumulative clustering of moonlighting proteins based on five features.

| Clusters | Sequences |
| --- | --- |
| Cluster-1 | {102, 103, 53, 20, 97, 119, 129, 164, 153, 120} |
| Cluster-2 | {79, 161} |
| Cluster-3 | {1, 5, 36, 93, 71, 104, 148, 18, 60, 37, 12, 21, 47, 163, 83, 127, 70, 84, 110, 22, 61, 90, 121, 134, 23, 94, 114, 128, 3, 88, 89, 137, 135, 136, 140} |
| Cluster-3 | {6, 165, 154, 112, 98, 25, 141, 33, 80, 118, 95, 8, 26, 155, 58, 51, 59, 145, 158, 31, 65, 123, 99, 44, 82, 48, 115, 96, 64, 10, 54, 117, 144, 159, 49, 78} |
| Cluster-3 | {106, 42, 116, 9, 109, 16, 45, 13, 113, 143, 152, 39, 43, 30, 124, 149, 28, 162, 2, 67, 57, 7, 62, 68, 38, 86, 138} |
| Cluster-4 | {11, 126, 50, 29, 81, 17, 73, 55, 101, 146, 147, 125, 24, 157, 91, 100, 41, 63, 27, 105, 52, 56, 32, 69, 107, 74, 75, 156, 160, 108, 131, 111} |
| Cluster-5 | {4, 77, 66, 85, 19, 35, 72, 151, 34} |
| Cluster-6 | {40, 142, 150, 133, 46} |

The distance matrix highlighted certain sequences. Specifically, sequences P09429 (Serial No. 79), Q99217 (Serial No. 87), P62328 (Serial No. 130), and P16402 (Serial No. 132) were each positioned more than 250 units apart from 119, 120, 155, and 90 other sequences, respectively. A closer examination of the mutual distances between these four sequences showed that P09429 was situated within 250 units of both P62328 and P16402. Notably, the most substantial observed distance of 416.895 was observed between sequences Q99714 and P62328.

### 4.8. Agglomerated distal-proximal relationships of moonlighting proteins

We identified a total of 19 moonlighting proteins as outliers based on the analysis of various extracted features, as detailed in previous sections (see Table 12). Within this group of 19 moonlighting proteins, three proteins namely Q99217 (Amelogenin), P62328 (Thymosin beta-4), and P16402 (Histone H1.3) stood out as outliers (Table 12) in cumulative clustering also. These three proteins exhibited distinctive roles in diverse biological processes, showcasing their moonlighting functions:

Q99217 plays a significant role in maintaining mitochondrial DNA.
P62328 functions as a secreted chemotaxis ligand and regulates the activation of actin polymerization and ILK kinase.
P16402 serves as a receptor for thyroglobulin.

**Table 12:** List of moonlighting proteins which turned out to be outliered based on various features namely 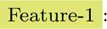 Relative frequency of amino acids; 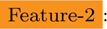 Homogeneous poly-string frequency; 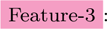 Relative frequency of four changes based on polar and nonpolar residues; 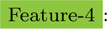 Relative frequency of nine changes based on acidic, basic and neutral residues 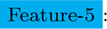 Relative frequency of sixteen changes based on disordered, highly flexible, moderately flexible, and other residues; 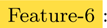 Structural and physicochemical features

| Serial Number | Uniprot ID | With respect to |
| --- | --- | --- |
| 21 | Q9P2J5 | Feature-2 |
| 27 | Q16637 | Feature-2 |
| 39 | P46060 | Feature-2 |
| 40 | P49790 | Feature-2, Feature-5 |
| 46 | O95997 | Feature-5 |
| 58 | P36897 | Feature-2 |
| 75 | O15105 | Feature-2 |
| 79 | P09429 | Feature-3, Feature-4, Feature-5 |
| 87 | Q99217 | Feature-1, Feature-4, Feature-6, Cumulative |
| 92 | P61254 | Feature-3 |
| 120 | P49759 | Feature-3 |
| 126 | P49792 | Feature-2 |
| 129 | Q5BJF6 | Feature-5 |
| 130 | P62328 | Feature-3, Feature-4, Feature-5, Feature-6, Cumulative |
| 132 | P16402 | Feature-1, Feature-5, Cumulative |
| 133 | P37198 | Feature-4 |
| 142 | Q96HA1 | Feature-4, Feature-5 |
| 150 | Q9Y6I3 | Feature-5 |
| 161 | P19338 | Feature-2, Feature-5 |

These 19 moonlighting proteins exhibited significant separation from the rest of the moonlighting proteins, as indicated by distinctive signature features originating from their amino acid sequences. It’s important to note that this separation does not necessarily correlate with their distances based on moonlighting function, as moonlighting functionality is typically independent of the protein’s primary structure [58].

Each of the six disjoint clusters as formed in cumulative clustering (Table 11) was further explored in detail to understand the proximal relationship among sequences of that cluster with respect to each feature individually. There may be more than one proximal set for a single feature and the sets were invariably disjoint. Proximal sets based on different features may or may not be disjoint. For example a subset of sequences belonging to cluster-1 formed one proximal set with respect to Feature-1 while producing three proximal sets with respect to Feature-5 (Table13). Further, sequence numbers 79 and 161 of cluster-2 set up a proximal set only for feature 5 and not for any other feature. Thus Table13 revealed how one or more than one proximal sets of moonlighting proteins with respect to individual feature finally contributed to the cumulative clustering. It is crucial to emphasize that the proximity of these sequential features does not necessarily imply a correlation with the moonlighting functions of the proteins. Following we present four types of examples of moonlighting proteins illustrating four possible different scenarios namely

- **Case-I:** Shared no proximal relationship based on sequence-features, but having identical moonlighting functions.
- **Case-II:** Shared proximal relationship based on sequence-features, but having distinct moonlighting functions.
- **Case-III:** Shared proximal relationship based on sequence-features along with identical moonlighting functions.
- **Case-IV:** Shared no proximal relationship based on sequence-features along with distinct moonlighting functions.

**Table 13:** Proximal sets of moonlighting proteins.

| Cluster-1: {102, 103, 53, 20, 97, 119, 129, 164, 153, 120} |  |
| --- | --- |
| Serial numbers | With respect to |
| {102, 103, 53, 20, 119, 129, 164, 153} | Feature-1 |
| {102, 103}, {53, 20, 97, 120}, {119, 164, 153} | Feature-5 |
| {102, 103, 129, 164, 153, 120} | Feature-4 |
| {102, 103} | Feature-6 |
| {102, 103, 53, 97, 119, 129, 164, 153} | Feature-3 |
| Cluster-2: {79, 161} |  |
| Serial numbers | With respect to |
| {79, 161} | Feature-5 |
| Cluster-3: {1, 5, 36, 93, 71, 104, 148, 18, 60, 37, 12, 21, 47, 163, 83, 127, 70, 84, 110, 22, 61, 90, 121, 134, 23, 94, 114, 128, 3, 88, 89, 137, 135, 136, 140, 6, 165, 154, 112, 98, 25, 141, 33, 80, 118, 95, 8, 26, 155, 58, 51, 59, 145, 158, 31, 65, 123, 99, 44, 82, 48, 115, 96, 64, 10, 54, 117, 144, 159, 49, 78, 106, 42, 116, 9, 109, 16, 45, 13, 113, 143, 152, 39, 43, 30, 124, 149, 28, 162, 2, 67, 57, 7, 62, 68, 38, 86, 138} |  |
| Serial numbers | With respect to |
| {1, 5, 36, 93, 71, 104, 148, 18, 60, 37, 12, 21, 47, 163, 83, 127, 70, 84, 110, 22, 61, 90, 121, 134, 23, 94, 114, 128, 3, 88, 89, 141, 8, 26, 155, 58, 51, 59, 145, 158, 65, 123, 99, 44, 82, 48, 96, 64, 117, 144, 159, 109, 16, 124, 28} | Feature-1 |
| {137, 135, 136, 140, 6, 165, 154, 112, 98, 25, 33, 80, 118, 95, 31, 115, 54, 78, 152} | Feature-1 |
| {49, 106, 9} | Feature-1 |
| {42, 116, 57, 7, 62, 68} | Feature-1 |
| {2, 67} | Feature-1 |
| {1, 5, 93, 148, 37, 12, 21, 47, 163, 83, 127, 110, 22, 61, 90, 114, 3, 88, 89, 137, 135, 136, 140, 49, 78, 106, 16, 149, 28} | Feature-4 |
| {36, 71, 104, 18, 60, 70, 84, 121, 134, 128, 165, 154, 112, 98, 25, 141, 33, 80, 118, 95, 8, 26, 155, 58, 51, 59, 158, 31, 123, 99, 44, 82, 115, 96, 64, 54, 117, 42, 116, 9, 109, 45, 13, 113, 143, 43, 162} | Feature-4 |
| {23, 94, 6, 145, 65, 48, 10, 144, 159, 152, 30, 124, 2, 67, 57, 7, 62, 68} | Feature-4 |
| {1, 5, 71, 104, 148, 18, 37, 21, 61, 90, 121, 23, 114, 3, 88, 89, 137, 135, 136, 140, 6, 165, 112, 98, 25, 33, 80, 118, 8, 26, 155, 58, 51, 59, 158, 65, 99, 115, 64, 49, 106, 42, 116, 45, 113, 162, 67, 57, 86, 138} | Feature-5 |
| {36, 93, 94, 141, 95, 145, 10, 54, 159, 9, 109, 16, 30, 2, 38} | Feature-5 |
| {60, 12, 47, 163, 83, 127, 22, 134, 44, 82, 13, 39, 124, 149, 7} | Feature-5 |
| {70, 84, 110, 128, 154, 31, 123, 96, 143, 152} | Feature-5 |
| {117, 144, 62} | Feature-5 |
| {5, 36, 93, 104, 60, 12, 21, 47, 163, 87, 127, 84, 110, 89, 141, 145, 158, 31, 65, 123, 99, 44, 82, 48, 96, 54, 117, 144, 62} | Feature-6 |
| {70, 23, 94} | Feature-6 |
| {135, 136} | Feature-6 |
| {140, 118, 78} | Feature-6 |
| {98, 151} | Feature-6 |
| {165, 154, 112, 39, 7} | Feature-6 |
| {10, 68} | Feature-6 |
| {5, 114, 128, 89, 137, 80, 96, 16, 162, 38, 86} | Feature-3 |
| {36, 93, 71, 104, 148, 18, 60, 37, 12, 21, 47, 163, 83, 127, 70, 84, 110, 22, 61, 90, 23, 3, 88, 8, 155, 31, 123, 45, 13, 149} | Feature-3 |
| {135, 136, 140, 6, 165, 154, 112, 25, 141, 33, 118, 95, 26, 58, 51, 59, 145, 158, 65, 99, 44, 82, 48, 115, 96, 64, 10, 54, 117, 144, 159, 49, 78, 106, 9, 109, 113, 143, 152, 39, 43, 30, 124, 28, 138} | Feature-3 |
| {98, 42, 116, 67} | Feature-3 |
| {2, 57, 7, 68} | Feature-3 |

*Example 1* (**Case-I**): Two proteins, P36897 (TGF-beta receptor type-1, Serial Number: 58) and Q99988 (Growth/ differentiation factor 15, Serial Number: 107), are not proximal in any of the five sequence-features. P36897 and Q99988 belonged to cluster-3 and cluster-4 respectively in cumulative clustering (Table 11). What makes this particularly intriguing is that both of these proteins share an identical moonlighting function, namely, “Nucleus/ Gene regulation.”

*Example 2* (**Case-II**): Two proteins, namely P08238 (Heat shock protein HSP 90-beta, Serial Number: 103) and P07900 (Heat shock protein HSP 90-alpha, Serial Number: 102), were observed to be in close proximity based on all five features. However, it’s noteworthy that despite their sequence-based similarity, these two proteins have distinct moonlighting functions. Specifically, HSP 90-alpha serves as a chaperone, contributing to MMP2 activation and wound healing through its involvement in exosomes, while HSP 90-beta plays a pivotal role in regulating the transcription machinery.

*Example 3* (**Case-III**): Our sequence-based phylogenetic analysis revealed that two moonlighting proteins, P02511 (alpha-crystallin B chain, Serial Number: 52) and P02489 (alpha-crystallin A chain, Serial Number: 56), share the same cluster (Cluster-4, as shown in Table 11). These proteins exhibit identical moonlighting functions as “Heat-shock proteins.”

*Example 4* (**Case-IV**): Q99217 (Amelogenin, Serial Number: 87) and P62328 (Thymosin Beta-4 (sequester actin), Serial Number: 130) turned out to be distal based on cumulative sequential features, and these two proteins have different moonlighting functions.

**Table 14:**
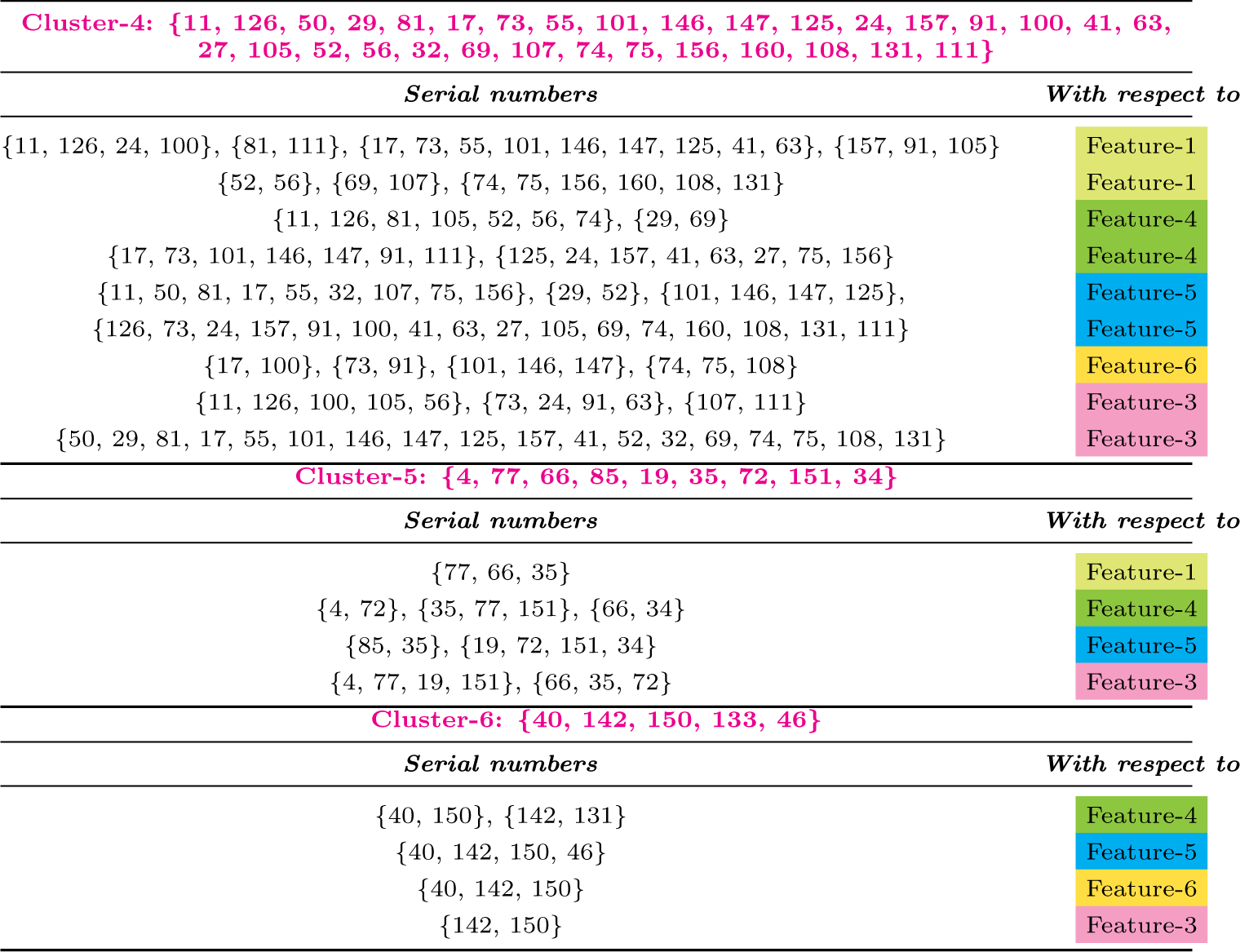
Proximal sets of moonlighting proteins.

| **Cluster-4: {11, 126, 50, 29, 81, 17, 73, 55, 101, 146, 147, 125, 24, 157, 91, 100, 41, 63, 27, 105, 52, 56, 32, 69, 107, 74, 75, 156, 160, 108, 131, 111}** |  |
| --- | --- |
| *Serial numbers* | *With respect to* |
| {11, 126, 24, 100}, {81, 111}, {17, 73, 55, 101, 146, 147, 125, 41, 63}, {157, 91, 105} | Feature-1 |
| {52, 56}, {69, 107}, {74, 75, 156, 160, 108, 131} | Feature-1 |
| {11, 126, 81, 105, 52, 56, 74}, {29, 69} | Feature-4 |
| {17, 73, 101, 146, 147, 91, 111}, {125, 24, 157, 41, 63, 27, 75, 156} | Feature-4 |
| {11, 50, 81, 17, 55, 32, 107, 75, 156}, {29, 52}, {101, 146, 147, 125}, | Feature-5 |
| {126, 73, 24, 157, 91, 100, 41, 63, 27, 105, 69, 74, 160, 108, 131, 111} | Feature-5 |
| {17, 100}, {73, 91}, {101, 146, 147}, {74, 75, 108} | Feature-6 |
| {11, 126, 100, 105, 56}, {73, 24, 91, 63}, {107, 111} | Feature-3 |
| {50, 29, 81, 17, 55, 101, 146, 147, 125, 157, 41, 52, 32, 69, 74, 75, 108, 131} | Feature-3 |
| **Cluster-5: {4, 77, 66, 85, 19, 35, 72, 151, 34}** |  |
| *Serial numbers* | *With respect to* |
| {77, 66, 35} | Feature-1 |
| {4, 72}, {35, 77, 151}, {66, 34} | Feature-4 |
| {85, 35}, {19, 72, 151, 34} | Feature-5 |
| {4, 77, 19, 151}, {66, 35, 72} | Feature-3 |
| **Cluster-6: {40, 142, 150, 133, 46}** |  |
| *Serial numbers* | *With respect to* |
| {40, 150}, {142, 131} | Feature-4 |
| {40, 142, 150, 46} | Feature-5 |
| {40, 142, 150} | Feature-6 |
| {142, 150} | Feature-3 |

### 4.9. Discussion

Moonlighting proteins, a unique class of multifunctional proteins, are characterized by a single polypeptide chain undertaking multiple physiologically relevant biochemical or biophysical roles. Moonlighting proteins are present in almost every organism that has been studied so far. Bacteria have approximetly 20 moonlighting proteins and humans have few hundred moonlighting proteins. These proteins are widely available in the entire evolutionary trees. Due to epigenetic changes, the activity of moonlighting proteins changes and they are evolutionary hotspots - as they are the first set of proteins which can sense the epigenetic changes. One more interesting point about moonlighting proteins is that each organism in the evolutionary tree has their own set of moonlight proteins. Therefore, Darwinian selection pressure has been employed-thus making moonlight proteins as checkpoints. One-hundred and sixty-five human moonlighting proteins have been obtained from the database MultitaskProtDBII, spanning a wide array of types, including receptors, enzymes, transcription factors, adhesins, and scaffolds. Remarkably, these proteins exhibit diverse combinations of canonical functions. While some of these proteins can execute both functions simultaneously, others adapt their role in response to environmental changes. The wide-ranging examples of identified moonlighting proteins and their potential benefits to organisms, such as orchestrating cellular activities, strongly suggest that many more moonlighting proteins remain to be unveiled. Furthermore, it was reported in the literature that amino acid sequence homology of moonlighting proteins may not necessarily possess moonlighting functions themselves [2, 8]. Therefore, relying solely on sequence homology might not be adequate for accurately predicting protein function. In this work, we investigate 165 human moonlighting proteins to discover various signature-features, which led us to develop a phylogenetic relationship among the set of moonlighting proteins. Signature-features include frequency of changes as obtained from change response profile based on three features, namely polar-nonpolar, acidic-badic-neutral, and intrinsic protein disorder. Phylogenetic analyses revealed a set of 19 moonlighting proteins having a distal relationship from remaining moonlighting proteins as obtained from different features (Table 12)). In the cumulative clustering, 156 moonlighting proteins formed six different clusters (Table 11) which were further analyzed in terms of proximity among moonlighting proteins for each sequence based feature. This distal-proximal relationship among these moonlighting proteins draws a distinct characterization of human moonlighting proteins, using unique genomic imprints that define these proteins within the human proteome. Furthermore, distal-proximal relationships might provide insights into the divergence and convergence of moonlighting protein functions.

This investigation sheds light on unique systems biology properties of the proteins and their association with certain physiological pathways especially for the characterization of the human multi-domain proteins. The study uncovers intriguing questions and hypotheses regarding the functional diversity of moonlighting proteins. These could serve as the basis for future research endeavors in the fields of moonlighting proteomics.

## Acknowledgements

Authors are very grateful to the laboratory supporters Mr. Ashim Dhar and Mrs. Piyali Maitra for their logistic assistance. Especially, authors would like to thank wholeheartedly Mr. Arindam Samanta for his insightful comments. Authors would like to thank the Department of Biotechnology, Govt. of India (BT/PR26647/NNT/28/1365/2017), Indian Space Research Organization (ISRO/RES/3/827/19-20), and Indian Statistical Institute (ISI) (ISI/TAC/PROJECT-1/2023-24) for their financial support.

## Author contributions statement

SSH, DN, AG, and VNU conceived the problem and theoretical experiments. DN, SSH, MS, VNU, Ankita Ghosh, and GD, and KL executed the results and performed the analyses. DN, SSH, VNU, MS, Ankita Ghosh, KL, and PB wrote the initial draft. All authors reviewed and edited the manuscript. SSH and AG supervised the entire project. All the authors checked, reviewed, and approved the final version of the manuscript.

---

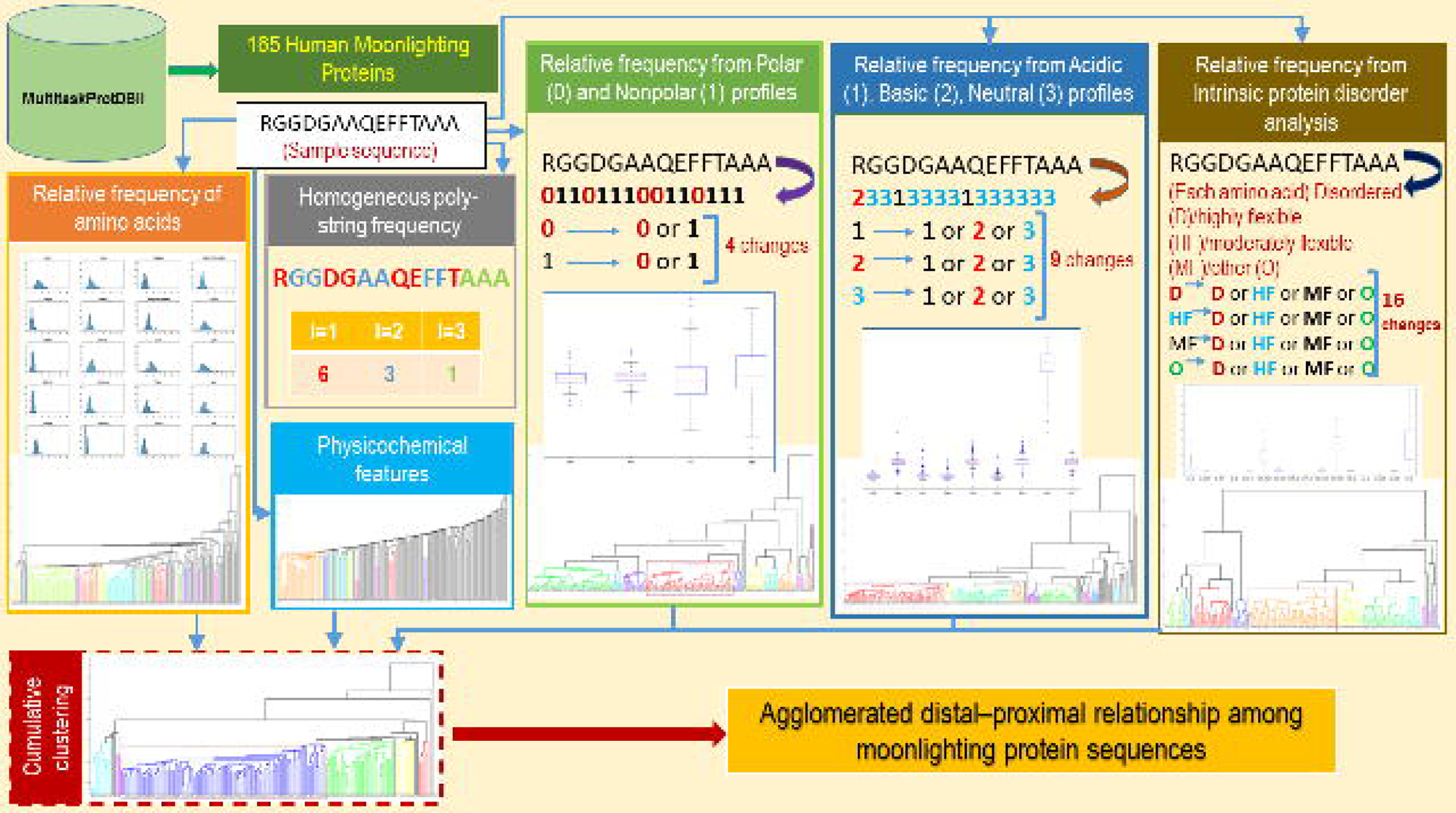

